# TADs pair homologous chromosomes to promote interchromosomal gene regulation

**DOI:** 10.1101/445627

**Authors:** Kayla Viets, Michael Sauria, Chaim Chernoff, Caitlin Anderson, Sang Tran, Abigail Dove, Raghav Goyal, Lukas Voortman, Andrew Gordus, James Taylor, Robert J. Johnston

## Abstract

Homologous chromosomes colocalize to regulate gene expression in processes including genomic imprinting and X-inactivation, but the mechanisms driving these interactions are poorly understood. In *Drosophila*, homologous chromosomes pair throughout development, promoting an interchromosomal gene regulatory mechanism called transvection. Despite over a century of study, the molecular features that facilitate chromosome-wide pairing are unknown. The “button” model of pairing proposes that specific regions along chromosomes pair with a higher affinity than their surrounding regions, but only a handful of DNA elements that drive homologous pairing between chromosomes have been described. Here, we identify button loci interspersed across the fly genome that have the ability to pair with their homologous sequences. Buttons are characterized by topologically associated domains (TADs), which drive pairing with their endogenous loci from multiple locations in the genome. Fragments of TADs do not pair, suggesting a model in which combinations of elements interspersed along the full length of a TAD are required for pairing. Though DNA-binding insulator proteins are not associated with pairing, buttons are enriched for insulator cofactors, suggesting that these proteins may mediate higher order interactions between homologous TADs. Using a TAD spanning the *spinelessd* gene as a paradigm, we find that pairing is necessary but not sufficient for transvection. *spineless* pairing and transvection are cell-type-specific, suggesting that local buttoning and unbuttoning regulates transvection efficiency between cell types. Together, our data support a model in which specialized TADs button homologous chromosomes together to facilitate cell-type-specific interchromosomal gene regulation.

## Introduction

Chromosomes are organized in a complex manner in the nucleus. In higher organisms, they localize to distinct territories (Cremer and Cremer, 2010). Regions of chromosomes interact to form compartments, which are segregated based on gene expression states (Eagen, 2018). Chromosomes are further organized into TADs, regions of self-association that are hypothesized to isolate genes into regulatory domains and ensure their activation by the correct *cis*-regulatory elements (Eagen, 2018). TADs vary in size from ~100 kb in *Drosophila melanogaster* to ~1 Mb in mammals *(Eagen, 2018; Sexton et al., 2012)*. Disruptions of nuclear organization, such as alteration of TAD structure and localization of genes to incorrect nuclear compartments, have major effects on gene expression (Clowney et al., 2012; Guo et al., 2015; Lupianez et al., 2015; Reddy et al., 2008). However, it is unclear how elements within the genome interact between chromosomes to organize chromatin and regulate gene expression.

One key aspect of nuclear architecture involves the tight colocalization, or “pairing,” of homologous chromosomes to facilitate regulatory interactions between different alleles of the same gene (Joyce et al., 2016). In *Drosophila melanogaster*, homologous chromosomes are paired throughout interphase in somatic cells (Stevens, 1906). This stable pairing provides an excellent paradigm to study the mechanisms driving interactions between chromosomes.

Despite over a century of study, it is poorly understood how homologous chromosomes come into close physical proximity. Classical studies of pairing proposed a “zipper” model, in which all regions of the genome have an equal ability to pair based on sequence homology, and chromosome pairing begins at the centromere and proceeds distally to the telomeres (Gelbart, 1982; Lewis, 1954). Studies of chromosome pairing initiation during development led to a shift in thinking towards the “button” model, which proposes that regions of high pairing affinity are interspersed along chromosome arms and come together through a random walk to initiate pairing (**Fig. 1A**) (Fung et al., 1998; Gemkow et al., 1998; Hiraoka et al., 1993).

**Figure 1:**
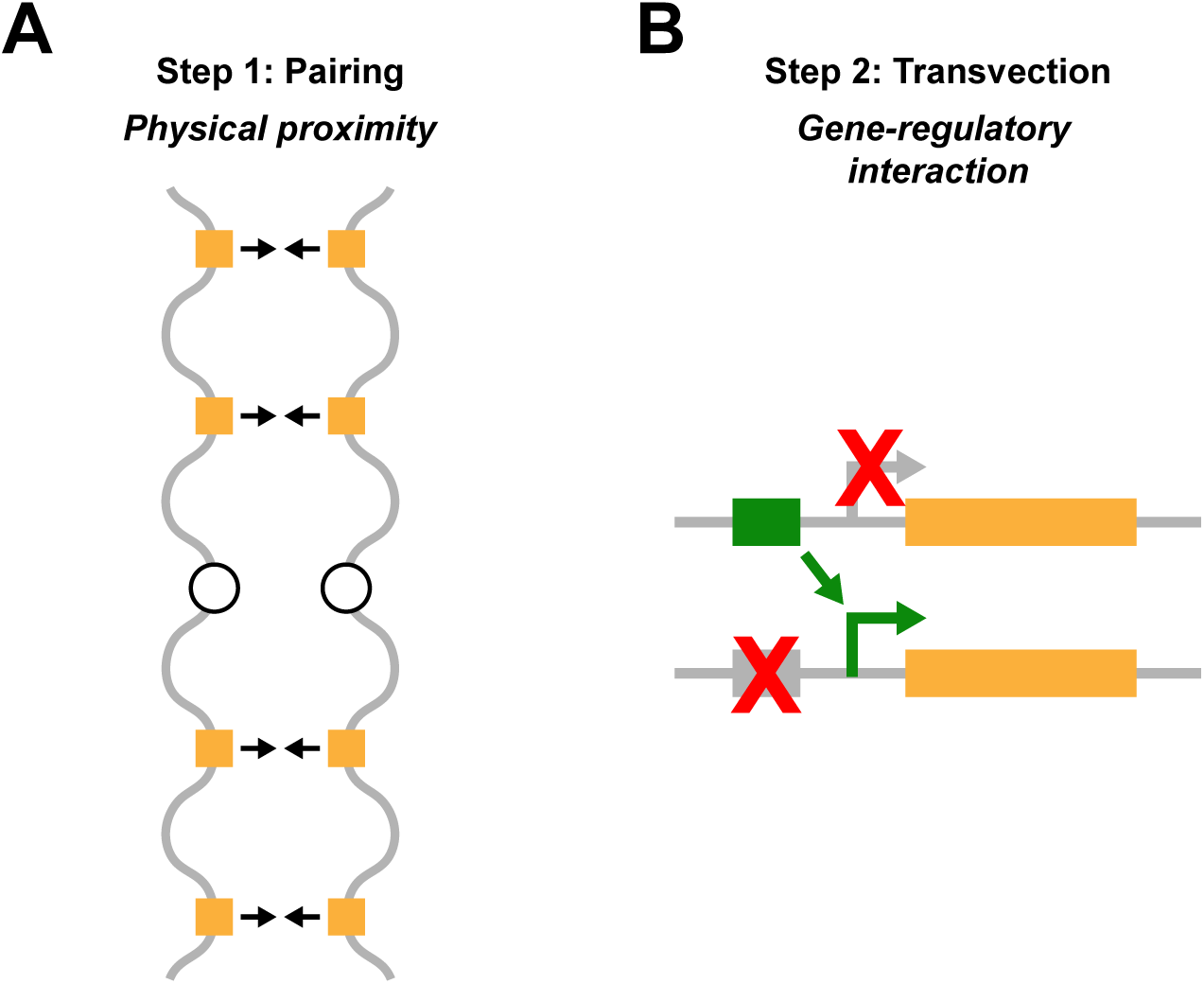
Homologous chromosomes “button” together to facilitate transvection. **A**. Button model of chromosome pairing. Yellow squares: button loci (high pairing affinity). **B**. During transvection, two different mutant alleles interact between chromosomes to rescue geneexpression. Green box: functional enhancer. Gray box with red X: mutated enhancer. Green arrow:functional promoter. Gray arrow with red X: mutated promoter.

The nature of the high-affinity “buttons” that bring homologous chromosomes together is still unclear. Many elements, including insulators, Polycomb Response Elements (PREs), and heterochromatic repeats, drive looping interactions in *cis* along the same chromosome arm or clustering between non-homologous sequences on different chromosomes (Blanton et al., 2003; Dernburg et al., 1996; Fujioka et al., 2016; Fujioka et al., 2009; Li et al., 2011; Li et al., 2013). However, only a handful of small DNA elements, including the gypsy retrotransposon and the Fab-7, Mcp, and TMR regions of the *Abd-B* locus, are known to drive pairing between identical homologous sequences on different chromosomes (Bantignies et al., 2003; Gerasimova et al., 2000; Li et al., 2011; Li et al., 2013; Ronshaugen and Levine, 2004; Vazquez et al., 2006). As three of these elements are within the same locus, the sequence and structural features that contribute to genome-wide homologous chromosome pairing are unknown. The scarcity of small DNA elements that are known to drive interchromosomal homologous pairing suggests that combinations of elements and/or higher order chromatin structures are required to button homologous chromosomes together.

Pairing of homologous chromosomes facilitates a gene-regulatory mechanism known as transvection, in which two different mutant alleles interact between chromosomes to rescue gene expression (**Fig. 1B**) (Lewis, 1954). Transvection has been described for a number of *Drosophila* genes (Duncan, 2002). With the exceptions of *Abd-B* and certain transgenes containing the Homie, gypsy, Mcp, TMR, and Fab7 sequences, transvection requires homologous chromosome pairing and is disrupted by chromosome rearrangements (Bantignies et al., 2003; Duncan, 2002; Fujioka et al., 2016; Gemkow et al., 1998; Hendrickson and Sakonju, 1995; Kravchenko et al., 2005; Lewis, 1954; Li et al., 2011; Muller et al., 1999; Sigrist and Pirrotta, 1997; Sipos et al., 1998; Vazquez et al., 2006; Zhou et al., 1999). DNA elements such as insulators and PREs contribute to transvection and similar phenomena at many loci across the genome (Bantignies et al., 2003; Fauvarque and Dura, 1993; Fujioka et al., 1999; Fujioka et al., 2016; Fujioka et al., 2009; Gindhart and Kaufman, 1995; Kapoun and Kaufman, 1995; Kassis, 1994; Kassis et al., 1991; Kravchenko et al., 2005; Li et al., 2011; Muller et al., 1999; Shimell et al., 2000; Sigrist and Pirrotta, 1997; Vazquez et al., 2006; Zhou et al., 1999), but it is unclear if the same DNA elements are always involved in both homologous chromosome pairing and transvection or if pairing and transvection are mechanistically separable.

Homologous chromosome pairing occurs more strongly in some cell types than in others. Pairing occurs in 15-30% of nuclei in the early embryo, gradually increases throughout embryonic development, and reaches a peak of 90-95% by the third instar larval stage (Dernburg et al., 1996; Fung et al., 1998; Gemkow et al., 1998; Hiraoka et al., 1993). Similarly, transvection efficiency varies widely between cell types (Bateman et al., 2012; Blick et al., 2016; Kassis et al., 1991; Mellert and Truman, 2012). However, a direct link between the level of pairing in a given cell type and the strength of transvection in that cell type has not been established.

Here, we develop a method to screen for DNA elements that pair and identify multiple button sites interspersed across the *Drosophila* genome, allowing us to examine the features that determine button activity. We find that a subset of TADs are associated with buttons and can drive pairing from different positions in the genome. By testing mutant alleles and transgenes of the *spineless* gene, we find that pairing and transvection are mechanistically separable and cell-type-specific. Together, our data suggest that TADs are one key feature of buttons that drive homologous chromosome pairing to promote cell-type-specific interchromosomal gene regulation.

## Results

### Identification of new button loci interspersed across the fly genome

Only a few elements are known to drive homologous pairing, and a majority of these elements are located within the *Abd-B* locus (Bantignies et al., 2003; Gerasimova et al., 2000; Li et al., 2011; Li et al., 2013; Ronshaugen and Levine, 2004; Vazquez et al., 2006), limiting the identification of general features that drive pairing throughout the genome. To look more broadly for elements that bring homologous chromosomes together, we selected transgenes from multiple locations in the genome, inserted them on heterologous chromosomes, and tested if they drove pairing with their endogenous loci. This assay provided a sensitized system for identifying regions of the genome that have an especially high affinity for homologous sequences, since it detected DNA sequences that pair outside of the context of chromosome-wide pairing.

We first screened a set of ~80-110 kb transgenes tiling a ~1 Mb region on chromosome 3R (**Fig. 2E**). We inserted individual transgenes into a site on chromosome 2L (site 1; **Fig. 2A**) and visualized their nuclear position using Oligopaints DNA FISH (Beliveau et al., 2012). As the endogenous and transgenic sequences were identical, we distinguished between them by labeling the sequence neighboring the endogenous locus with red probes and the sequence neighboring the transgene insertion site with green probes (“2 color strategy”)(**Fig. 2A**). We examined pairing in post-mitotic larval photoreceptors to avoid disruptions caused by cell division.

**Figure 2:**
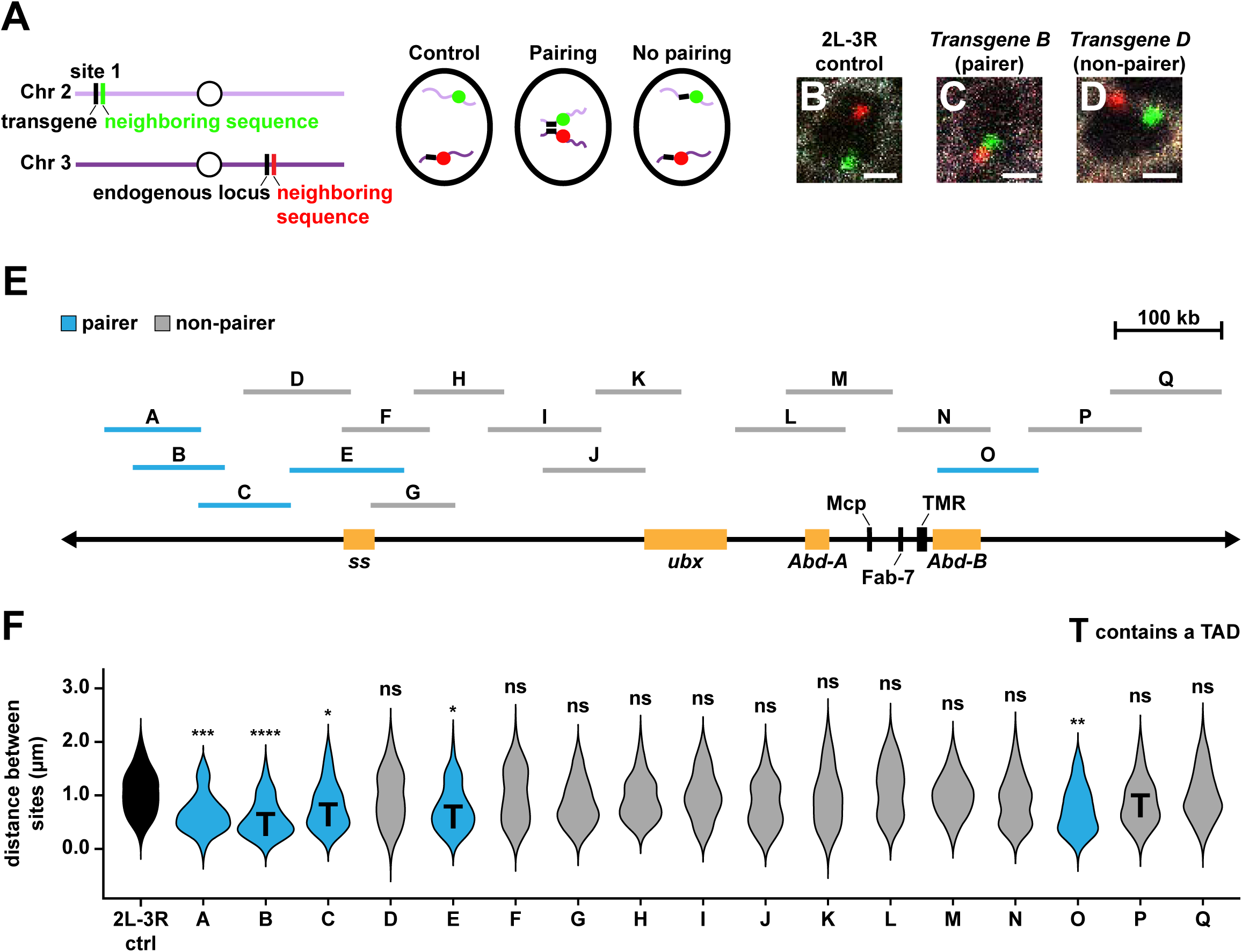
A screen for pairing elements identifies buttons interspersed along chromosome 3R. **A**. Two-color DNA FISH strategy. In controls, red and green FISH punctae on heterologous chromosomes are far apart in the nucleus. If a transgene drives pairing, red and green FISH punctae are close together in the nucleus. If a transgene does not drive pairing, red and green FISH punctae are far apart in the nucleus, similar to a control. **B-D**. Control, pairer, and non-pairer examples. Image for 2L-3R control is from the same experiment as in **Fig. 3B, 5G**, and **S16A**. Scale bars=1 μm. White: Lamin B, red: probes neighboring endogenous sequence, green: probes neighboring transgene insertion site. **E**. ~1 Mb region of chromosome 3R used for pairing screen. Orange boxes indicate locations of themajor developmental genes *spineless* (*ss*), *ultrabithorax* (*ubx*), *Abdominal-A* (*Abd-A*), and *Abdominal-B* (*Abd-B*). Black lines indicate the locations of the pairing elements Mcp, Fab-7, and TMR. **F**. Quantifications for all transgenes from the initial screen. Black: control, blue: pairers, gray: non-pairers. **T:** contains a TAD. Control data are the same as in **Fig. 3E** (2L-3R control), **3I**, **6C** (2L-3Reye control), **S1A-B, S12J**, and **S14Q** (2L-3R control). *Transgene D-G* data are the same as in **Fig.3I** and **S1B**. *Transgene H-K, M, P, Q* data are the same as in **Fig. S1B**. ****=p<0.0001, ***=p<0.001,**=p<0.005, *=p<0.05, ns=p>0.05, one-way ANOVA on ranks with Dunn’s multiple comparisons test.

To determine whether a transgene drove pairing, we compared the 3-D distance between the transgene and its endogenous site to the 3-D distance of a negative control. If the distance between the insertion site and the endogenous site was significantly lower with a transgene present than in the negative control, then the transgene drove pairing (**Fig. 2A**). The negative control measured the distance between the transgene insertion site on chromosome 2L and a site on chromosome 3R in a wild type background. Since different loci along a chromosome arm might vary in their distance from other chromosomes, we examined additional negative controls in multiple cell types and observed no statistical differences (**Fig. S1A**). Thus, for all transgene pairing experiments described in this paper, we used single negative controls for each chromosome arm tested.

To determine the statistical significance of transgene pairing relative to a control, we first tested each transgene dataset for a Gaussian distribution. In instances where transgene distance distributions were non-Gaussian, we tested for statistical significance by comparing the median of each transgene distribution to the negative control (**Fig. 2F**). Gaussian transgene distance distributions were tested for significance using a t-test to compare means. In all cases where more than one transgene was compared to a control, we corrected for multiple comparisons.

Only a subset (5/17) of transgenes (“pairers”) in our initial screen drove pairing between chromosomes 2L and 3R in our initial screen, bringing the distances between the red and green FISH signals significantly closer than in the negative control (**Fig. 2B-C, F; Fig. S2A**). The red and green signals did not completely overlap, likely because they did not directly label the paired sites (**Fig. S3A-C**). For the remaining 12/17 transgenes (“non-pairers”), the distances between the red and green signals were not significantly different from the negative control, indicating that they did not drive pairing (**Fig. 2B, D, F; Fig. S2B**).

To confirm our assignment of pairers vs. non-pairers, we used maximum likelihood estimation to fit our transgene data to single or double Gaussian distributions. 4/5 pairers fit a double Gaussian distribution, indicating a split between paired and unpaired populations of nuclei. The remaining pairer, *Transgene E*, fit a single Gaussian distribution, but the mean of the *Transgene E* distribution was significantly lower than the mean of the negative control distribution, indicating a high degree of pairing (**Fig. S1B**). Conversely, only 2/12 non-pairers (*Transgenes L* and *N*) fit a double Gaussian distribution. Since the median distances for *Transgenes L* and *N* were not significantly different from the negative control (**Fig. 2F**), these transgenes may drive “weak” pairing at a level significantly lower than the other pairers we identified. The remaining 10/12 non-pairers fit a single Gaussian distribution, and their means did not significantly differ from the negative control (**Fig. S1B**). Thus, our screen identified multiple new button elements.

The pairing observed between transgenes on chromosome 2L and endogenous sequences on chromosome 3R could be affected by the transgene insertion site. To test the position independence of button pairing, we inserted *Transgene E* onto chromosome 3L (site 3; **Fig. S4A**) and found that it paired with its homologous endogenous locus on chromosome 3R (**Fig. S4B-D**), showing that buttons can drive pairing from different sites in the genome.

Thus, we identified multiple button loci along chromosome 3R that overcome endogenous nuclear architecture to drive pairing between non-homologous chromosomes.

### TADs are features of buttons

We next sought to determine common features between the new buttons we identified, focusing first on chromatin structure. We examined 14 publicly available Hi-C datasets to determine the relationship between buttons and topologically associated domains (TADs), genomic regions of self-association. We defined TADs using directionality indices, which measure the bias of a genomic region towards upstream or downstream interactions along the chromosome (Dixon et al., 2012). TADs on a directionality index are read from the beginning of a positive peak, which indicates downstream interactions, to the end of a negative peak, which indicates upstream interactions (**Fig. S5A; Fig. S6A-E; Fig. S7A-E**).

We found that 60% of pairers encompassed a complete TAD, including both TAD boundaries, compared to only 8% of non-pairers (**Fig. 2F; Fig. S5A; Fig. S8A-B**), suggesting that specific TADs contribute to button function. To test the hypothesis that TADs are a feature of buttons, we selected six transgenes encompassing entire TADs on chromosomes X, 2L, 2R, and 3R (**Fig. 3E; Fig. S6A-E; Fig. S7C; Fig. S8A**) and compared them to four transgenes that did not encompass entire TADs, taken from chromosomes X, 2L, and 3R (**Fig. 3E; Fig. S2E; Fig. S7A-B, D-E; Fig. S8B**). Based on the availability of Oligopaints probes, we used an alternative FISH strategy for a subset of these transgenes, in which the identical transgene and endogenous sequences were labeled with the same red fluorescent probes (**Fig. 3A**). With this 1-color strategy, FISH punctae ≤0.4 μm apart could not be distinguished as separate and were assigned a distance of 0.4 μm apart (see Materials and Methods).

**Figure 3:**
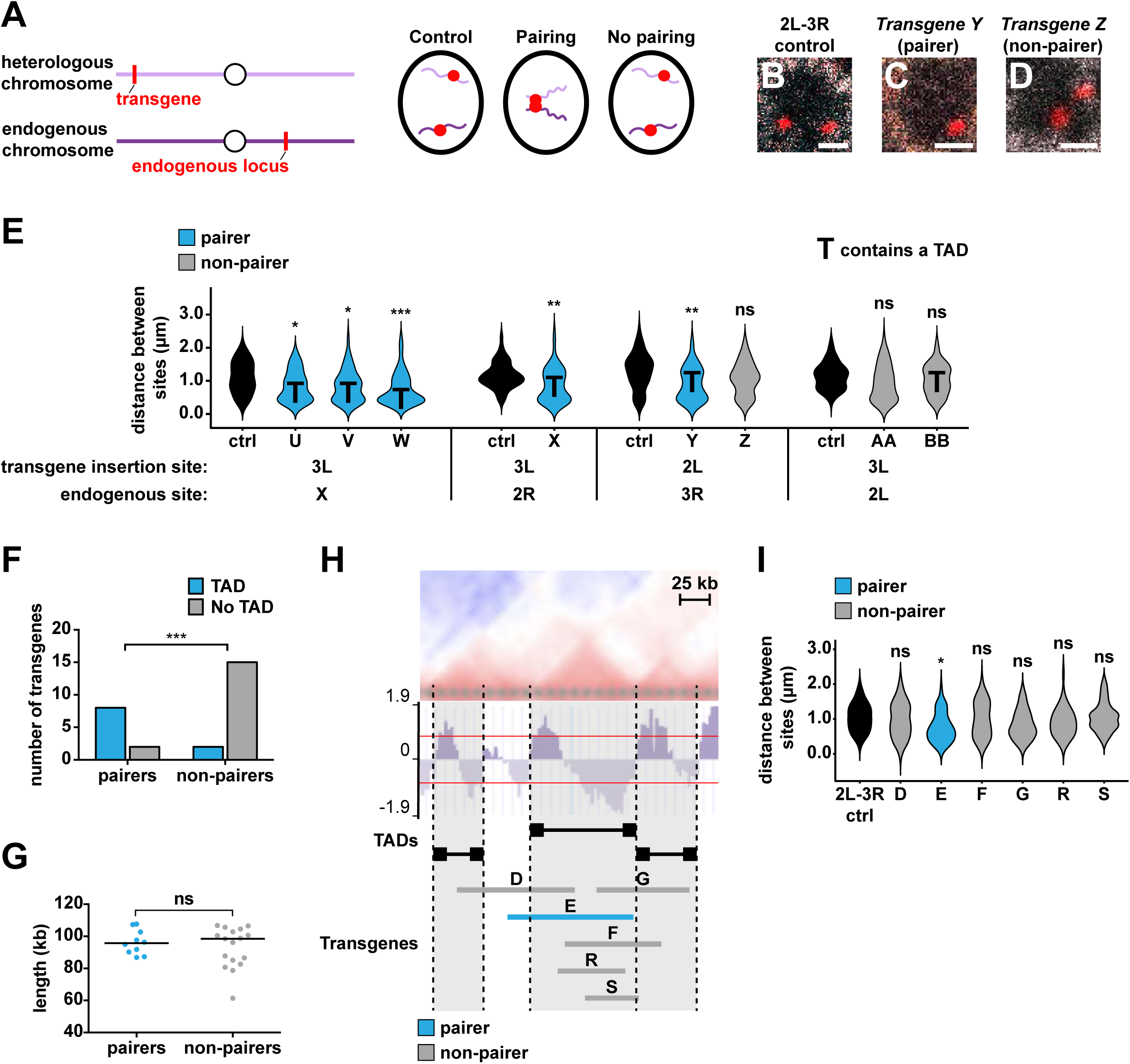
Specialized TADs contribute to button activity and drive pairing. **A**. One-color DNA FISH strategy: In controls, two red FISH punctae on heterologous chromosomes are far apart in the nucleus. If a transgene drives pairing, the two red FISH punctae are close together in the nucleus and indistinguishable as separate dots. If a transgene does not drive pairing, the two red FISH punctae are far apart in the nucleus, similar to a control. **B-D**. Control, pairer, and non-pairer examples. Scale bars=1 μm. White: Lamin B, red: probes against endogenous sequence and transgene. Image for 2L-3R control is from the same experiment as **Fig. 2B**, **5G**, and **S16A**. Image for *Transgene Z* is from the same experiment as **Fig. S3B**. **E**. Quantifications for additional transgenes. **T:** contains a TAD. Black: controls, blue: pairers, gray:non-pairers. ***=p<0.001, **=p<0.005, *=p<0.05, ns=p>0.05, one-way ANOVA on ranks with Dunn’smultiple comparisons test (for *Transgenes U-W*, *Y-Z*, *AA-BB*) or Wilcoxon rank-sum test (for*Transgene X*). 3L-X control data are the same as in **Fig. S1A-B** and **S2E**. 3L-2R control data and 3L-2L control data are the same as in **Fig. S1A**. 2L-3R control data are the same as in **Fig. 2F, 3I, 6C**(2L-3R eye control), **S1A-B, S12J**, and **S14Q** (2L-3R control). Controls were imaged in two colors,then pseudocolored red and re-scored in one color. **F**. Number of pairers vs. non-pairers spanning TADs. Blue: spans a TAD. Gray: does not span aTAD. ***=p<0.001, Fisher’s exact test. **G**. Comparison of length for all pairers vs. non-pairers tested in **Fig. 2F, 3E**, and **S2E**. Blue: pairers,gray: non-pairers. ns=p>0.05, Wilcoxon rank-sum test. Black lines indicate medians. **H**. Representative Hi-C heat map and directionality index (NCBI GSE38468) showing TADs in the region covered by *Transgenes D-G*. Dotted lines: TAD boundaries. Black bars: TADs. Red lines indicate a directionality index signal of 0.8 or −0.8, the cutoff for a TAD. See **Fig. S5A** for TAD assessment. **I**. Quantifications for transgenes that “split” the TAD covered by *Transgene E* in **Fig. 3H**. Black: control, blue: pairers, gray: non-pairers. *=p<0.05, ns=p>0.05, one-way ANOVA on ranks with Dunn’s multiple comparisons test. Control data are the same as in **Fig. 2F, 3E** (2L-3R control), **6C** (2L-3R eye control), **S1A-B, S12J**, and **S14Q** (2L-3R control). *Transgene D, F*, and *G* data are the same as in **Fig. 2F** and **S1B**. *Transgene E* data are the same as in **Fig. 2F** and **S1B**. *Transgene S* data are the same as in **Fig. S14Q** (*Transgene S* site 1).

5/6 transgenes that spanned entire TADs drove pairing, while 0/4 transgenes that did not span a TAD drove pairing (**Fig. 3B-E; Fig. S2C-E**), further supporting the importance of TADs for button activity. In total for all transgenes tested in **Fig. 2F**, **Fig. 3E**, and **Fig. S2E**, 80% of pairers spanned a TAD (8/10) while only 12% of non-pairers spanned a TAD (2/17) (**Fig 3F**), indicating that specific TADs contribute to button activity and drive pairing. Transgenes of near identical lengths had different pairing abilities, further suggesting that the content of a transgene (i.e. TADs) determines pairing (**Fig. 3G**).

The ~80-110 kb size limitation of publicly available transgenes prevented testing larger TADs for pairing with our transgene assay. Transgenes that covered only parts of a large TAD on chromosome 3R did not drive pairing (**Fig. S9A**). To test this large TAD for pairing, we utilized a 460kb duplication of chromosome 3R onto chromosome 2R (**Fig. S9B**), which encompassed the entire TAD (**Fig. S5A; Fig. S9A**). We found that the duplication drove pairing with its homologous endogenous site (**Fig. S9C-E**), further supporting a role for TADs in pairing.

Entire TADs could drive pairing, or smaller elements contained within TADs could bring homologous regions together. We therefore examined the effects of “splitting” a TAD, focusing on the TAD spanned by *Transgene E. Transgene D*, which covers the 5’ end of *Transgene E*, did not drive pairing (**Fig. 3H-I; Fig. S2B**). *Transgenes G* and *S*, which cover the 3’ end of *Transgene E*, also did not drive pairing (**Fig. 3H-I; Fig. S2B**). Because there is a ~19 kb gap between *Transgenes D* and *G*, we hypothesized that this 19 kb region might contain an element required for pairing. However, *Transgenes F* and *R*, which both contained this 19 kb region, did not drive pairing. Thus, on their own, the 5’, middle, or 3’ regions of the *Transgene E* TAD are not sufficient to drive pairing. Together, these observations suggest that a combination of elements interspersed across the complete *Transgene E* TAD drive pairing.

### Examining the relationship between pairing, chromatin factors, and gene expression state

We next examined other features that could contribute to homologous chromosome pairing. As insulators have been linked to long-distance chromosome interactions(Blanton et al., 2003; Dernburg et al., 1996; Fritsch et al., 2006; Fujioka et al., 2016; Fujioka et al., 2009; Li et al., 2011; Li et al., 2013), we tested whether the number of binding sites for individual *Drosophila* insulator proteins was higher in pairers than in non-pairers. Using previously published ChIP-chip and ChIP-seq data(Cuartero et al., 2014; Negre et al., 2010; Ong et al., 2013; Van Bortle et al., 2012; Wood et al., 2011), we found no association between pairing and any of the DNA-binding insulator proteins (BEAF, Su(Hw), CTCF, and GAF)(**Fig. 4A-D**), suggesting that complex combinations of bound insulators, rather than individual proteins, might contribute to homologous pairing or stabilize interactions between already paired TADs(Lim et al., 2018). Intriguingly, both Cp190 and Mod(mdg4), insulator co-factors that do not directly bind to DNA, were associated with pairing (**Fig. 4E-F**). These proteins may mediate interactions between complex clusters of DNA-binding insulator proteins or play additional roles independent of insulator function to bring homologous chromosomes together.

**Figure 4:**
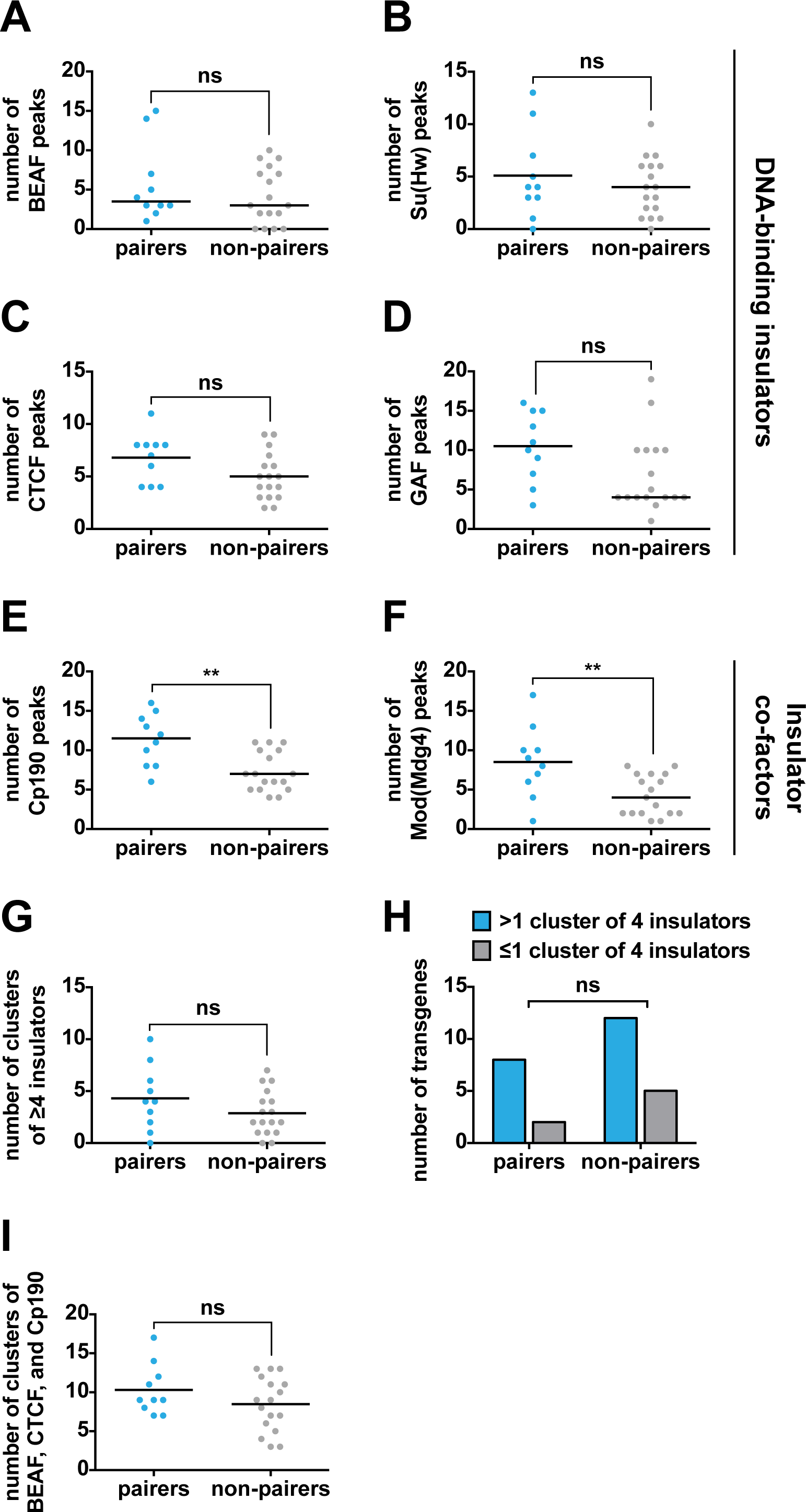
Examining the relationship between insulator binding sites and pairing. **A-I**. Quantifications for pairers and non-pairers tested in **Fig. 2F, 3E**, and **S2E**. **=p<0.01, ns=p>0.05, Wilcoxon rank-sum test (**A, D-F**), unpaired t-test with Welch’s correction (**B-C, G, I**), or Fisher’s exact test (**H**). The Wilcoxon rank-sum test compares medians, while the unpaired t-test with Welch’s correction compares means. Therefore, the black lines in **A, D-F** indicate medians, and the black lines in **B-C, G, I** indicate means. **A-G, I:** Blue: pairers, gray: non-pairers. **A**. BEAF ChIP peaks. **B**. Su(Hw) ChIP peaks. **C**. CTCF ChIP peaks. **D**. GAF ChIP peaks. **E**. Cp190 ChIP peaks. **F**. Mod(mdg4) ChIP peaks. **G**. Number of clusters of ≥4 insulators. **H**. Number of pairers vs. non-pairers with >1 or ≤1 cluster of 4 insulators. Blue: >1 cluster, gray: ≤1 cluster. **I**. Number of clusters containing any combination of BEAF, CTCF, and Cp190.

We next investigated the relationship between pairing and insulator clustering, defining a cluster as any region bound by ≥4 unique insulator proteins, and observed no association with pairing (**Fig. 4G-H**). As *Drosophila* TAD boundaries are enriched for CTCF, BEAF, and Cp190 (Sexton et al., 2012), we examined the association between pairers and clusters of these insulators. Again, we observed no relationship with pairing (**Fig. 4I**), suggesting that a complete TAD, rather than the insulators clustered at its boundaries, is needed for pairing. Thus, we do not find a strong relationship between insulators and pairing. However, it is possible that even larger clusters of insulators, in combination with other architectural proteins, mediate pairing between TADs, or that each button has a unique “code” of insulators bound across the entire TAD region that assist in driving pairing.

Because the small DNA elements Mcp, Fab-7, and TMR drive pairing between homologous chromosomes (Bantignies et al., 2003; Li et al., 2011; Li et al., 2013; Ronshaugen and Levine, 2004; Vazquez et al., 2006), we next examined whether these elements drove pairing in our assay. Unexpectedly, *Transgene M*, which contained Mcp, and *Transgene N*, which contained Fab-7 and TMR, were both non-pairers (**Fig. 2E-F; Fig. S2B**), suggesting that pairing driven by these elements may be context-specific, or that these elements drive pairing at a low level that cannot be distinguished by this assay.

To investigate additional elements that contribute to pairing, we examined modENCODE ChIP data and found no association between pairing and Polycomb Group (PcG) binding sites, repressing epigenetic marks, or non-coding RNAs (ncRNAs)(**Fig. S10A-F**).

Gene activity plays a critical role in nuclear architecture: individual TADs interact with each other in compartments, which are partitioned by expression state(Eagen, 2018). The pairing we observe between transgenes and their endogenous loci might simply be a result of segregation of the genome into active (A) and repressed (B) compartments. We hypothesized that if compartmentalization alone drives pairing, then the distance between any two active regions or any two repressed regions should be less than the distance between an active region and a repressed region. To test this hypothesis, we performed RNA-seq on larval eye discs, the tissue we used in our pairing experiments, to identify active or repressed regions. We selected two loci on different chromosomes that were highly expressed (A1 and A2; **Fig. S11A, C**), and two loci that were expressed at low levels (B1 and B2; **Fig. S11B-C**). The level of A1-A2 and B1-B2 interaction did not differ from a negative control or from the level of A1-B2 interactions (**Fig. S11D-H**). Moreover, we found no association between pairing and active transcription (**Fig. S11I-J**). Our data suggest that pairing is driven by specific interactions between homologous TADs, rather than general interactions based on expression state alone.

### Pairing and transvection occur despite chromosomal rearrangements

We next interrogated the relationship between pairing and the gene regulatory process of transvection. Chromosomal rearrangements have been shown to disrupt pairing of genes located near rearrangement breakpoints (Duncan, 2002; Lewis, 1954). However, we observed pairing of ~100 kb transgenes with their endogenous loci, suggesting that intact homologous chromosomes are not required for pairing and that pairing driven by TADs tolerates nearby breakpoints. Supporting our observations, the *Abd-B* locus pairs in the presence of chromosomal rearrangements (Gemkow et al., 1998; Hendrickson and Sakonju, 1995; Sipos et al., 1998). We therefore reexamined how rearrangements affect pairing, focusing on a button defined by a TAD spanning the *spineless (ss)* locus (“*ss* button”; **Fig. 5A**).

**Figure 5:**
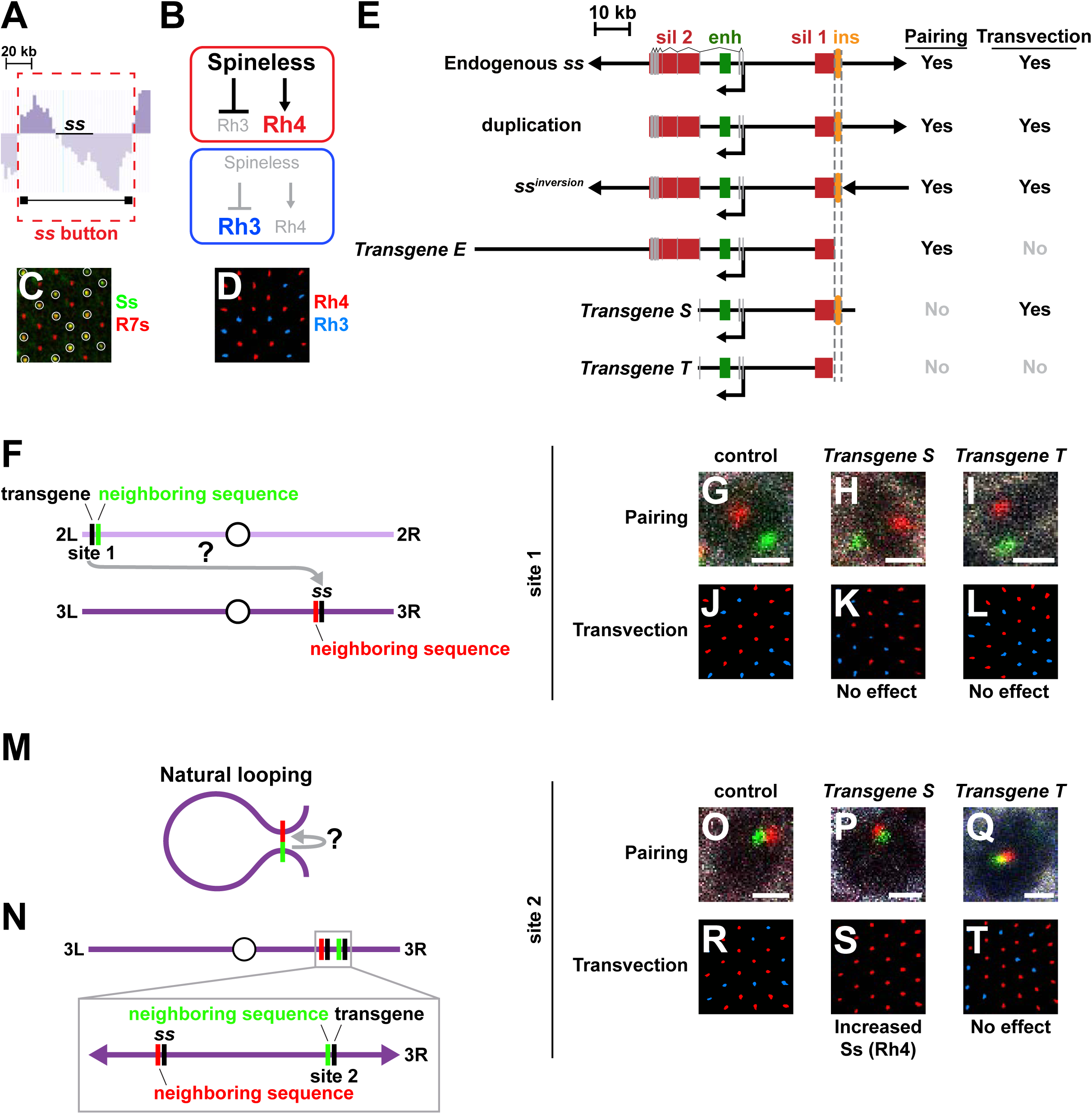
Pairing is necessary but not sufficient for transvection. **A**. Representative directionality index (NCBI GSE38468) showing the TAD that defines the *ss* button.Black bar: TAD. See **Fig. S5A** for TAD assessment. **B**. Spineless (Ss) activates Rh4 and represses Rh3. **C**. Ss is expressed in ~70% of R7s. Green: Ss, red: Prospero (R7 marker), white circles: Ss-expressing R7s. **D**. Rh3 (blue) and Rh4 (red) expression in wild type R7s. **E**. *ss* alleles and transgenes. ins: insulator, sil 1: silencer 1, enh: enhancer, sil 2: silencer 2. Smallerblack arrows: transcription start sites. Gray rectangles: exons. Dotted gray lines: region required fortransvection. **F**. Strategy used to assess pairing and transvection from site 1 in **Fig. 5G-L**. Gray arrow with “?”indicates that *Transgenes S* and *T* were tested for transvection. **G-I, O-Q**. Scale bars= 1 μm. White: Lamin B, red: probes neighboring endogenous sequence, green: probes neighboring transgene insertion site. **J-L, R-T**. Red: Rh4, blue: Rh3. **G-I**. Pairing assay images of *2L-3R control*, *Transgene S site 1*, and *Transgene T site 1*. See **Fig. S14Q** for quantifications. Image for 2L-3R control is from the same experiment as **Fig. 2B, 3B**, and **S16A**. **J-L**. Rh3 and Rh4 expression in *wild type control* (Ss(Rh4)=70%), *Transgene S site 1* (Ss(Rh4)=57%), and *Transgene T site 1* (Ss(Rh4)=55%). The slight decrease in Rh4 frequency for *Transgene S site 1* and *Transgene T site 1* is likely due to background genetic effects. **M**. Natural chromosome looping forces transgenes into close proximity with endogenous *ss*, mimicking pairing and facilitating transvection. Gray arrow with “?” indicates that *Transgenes S* and *T* were tested for transvection. **N**. Strategy used to assess pairing and transvection from site 2 in **Fig. 5O-T**. **O-Q**. Pairing assay images of *site 2 control*, *Transgene S site 2*, and *Transgene T site 2*. See **Fig. S14Q** for quantifications. **R-T**. Rh3 and Rh4 expression in *wild type control* (Ss(Rh4)=70%), *Transgene S site 2* (Ss(Rh4)=98%) and *Transgene T site 2* (Ss(Rh4)=78%).

To assess the effects of local rearrangements on *ss* button pairing, we examined a naturally occurring chromosomal inversion with a breakpoint immediately upstream of *ss* (*ss^inversion^*) and a duplication with a breakpoint immediately downstream of *ss* (**Fig. 5E**). Both *ss^inversion^* and the duplication paired with endogenous *ss* (**Fig. S9C-E; Fig. S12A-B**), showing that *ss* button pairing occurs despite chromosomal rearrangements. Consistent with these findings, pairing also occurred at the *ss* locus in flies with balancer chromosomes containing numerous large inversions and rearrangements (**Fig. S12F-J**). Thus, similar to *Abd-B*, pairing of *ss* occurs despite chromosomal rearrangements, consistent with a model in which homologous TADs find each other in the nucleus independent of chromosome-wide homology.

Pairing is required for the genetic phenomenon of transvection, in which DNA elements on a mutant allele of a gene act between chromosomes to rescue expression of a different mutant allele (**Fig. 1B**). In cases where chromosomal rearrangements perturb pairing, transvection is also disrupted (Duncan, 2002; Lewis, 1954). Since chromosomal rearrangements did not ablate pairing at the *ss* button, we hypothesized that transvection would occur at the *ss* locus in these genetic conditions.

In the fly eye, Ss is normally expressed in ~70% of R7 photoreceptors to activate expression of Rhodopsin 4 (Rh4) and repress Rhodopsin 3 (Rh3; **Fig. 5B-D**). Ss is absent in the remaining 30% of R7s, allowing Rh3 expression (**Fig. 5B-D**) (Wernet et al., 2006). Regulatory mutations in the *ss* gene cause decreases or increases in the ratio of Ss^ON^: Ss^OFF^ cells. When two *ss* alleles with different ratios are heterozygous, transvection between chromosomes (also known as Interchromosomal Communication) determines the final ratio of Ss^ON^: Ss^OFF^ R7s (Johnston and Desplan, 2014). Thus, the Ss^ON^: Ss^OFF^ ratio is a phenotype that allows for quantitative assessment of transvection. Throughout our *ss* transvection experiments, we evaluated Rh3 and Rh4 expression, as they faithfully report Ss expression in R7s (i.e. Ss^ON^ = Rh4; Ss^OFF^ = Rh3). We previously observed transvection at the *ss* locus for the duplication and balancer chromosome alleles (Johnston and Desplan, 2014). We similarly observed transvection at the *ss* locus for the *ss^inversion^* allele (**Fig. S12C-E**). Together, these data suggested that buttons can drive pairing and transvection despite chromosomal rearrangements.

### Pairing is necessary but not sufficient for transvection

As chromosomal rearrangements did not impair *ss* pairing or transvection, we further investigated the relationship between pairing and transvection using *ss* transgenes. Both *Transgene S* and *Transgene T* are expressed in 100% of R7s because they lack a silencer DNA element, but do not produce functional Ss protein because they lack critical coding exons (**Fig. 5E; Fig. S13A-J**)(Johnston and Desplan, 2014). *Transgene T* differs from *Transgene S* in that it lacks 6 kb at its 5’ end (**Fig. 5E**). We predicted that if *Transgenes S* and *T* performed transvection, they would upregulate expression of endogenous *ss*.

When inserted onto chromosomes 2L or 3L (sites 1 and 3; **Fig. 5F; Fig. S14A**), *Transgenes S* and *T* did not drive pairing with the endogenous *ss* locus on chromosome 3R (**Fig. 5G-I; Fig. S14B-C, E, Q**). At these sites, *Transgenes S* and *T* did not upregulate *ss* expression, indicating that they could not perform transvection when unpaired (**Fig. 5J-L; Fig. S14D, F**).

We next wondered whether *Transgenes S* and *T* could perform transvection if we mimicked pairing by forcing them into close physical proximity with endogenous *ss*. We performed a FISH screen to identify genomic sites that naturally loop to endogenous *ss* (**Fig. 5M**) and identified three such sites, located 4.8 Mb upstream of *ss*, 0.4 Mb upstream of *ss*, and 4.6 Mb downstream of *ss* (sites 2, 4, and 5; **Fig. 5M-O; Fig. S14G-H, M-N, Q**).

When we inserted *Transgene S* at these sites, it was forced into close proximity with endogenous *ss* (**Fig. 5P; Fig. S14I, O, Q**) and upregulated Ss (Rh4) into nearly 100% of R7s (**Fig. 5R-S; Fig. S14J, P**)(Johnston and Desplan, 2014). Thus, natural chromosome looping can force loci into proximity and, like pairing, facilitate transvection. In contrast, when we forced *Transgene T* into close proximity with endogenous *ss* (**Fig. 5Q; Fig. S14K, Q**), it did not upregulate Ss (Rh4) expression (**Fig. 5T; Fig. S14L**), indicating that it could not perform transvection even when paired. Thus, pairing is necessary but not sufficient for transvection.

We compared the DNA sequence of *Transgene T*, which does not perform transvection, to *Transgene S*, the *ss^inversion^*, and the duplication, which perform transvection. An upstream region of ~1.6 kb is present in *Transgene S*, the *ss*^*inversion*^, and the duplication, but missing from *Transgene T*, suggesting that this region contains a critical element for transvection (**Fig. 5E**). ModENCODE ChIP data showed that this region was bound by the *Drosophila* insulator proteins CTCF, BEAF, Mod(Mdg4), and Cp190. Additionally, this DNA sequence performed P-element homing (Johnston and Desplan, 2014), an indicator of insulator activity. Together, these data suggested that the DNA element required for transvection is an insulator.

To further test whether this insulator was required for transvection, we examined *Transgene E*, which drove pairing and contained the complete *ss* locus, except for the insulator element (**Fig. 2F; Fig. 5E; Fig. S2A; Fig. S4B-D**). We utilized genetic backgrounds in which *Transgene E* was the only source of Ss protein, so that any changes in Ss (Rh4) expression would indicate transvection effects on *Transgene E*. As a control, we examined *Transgene E* expression when the endogenous *ss* locus was hemizygous for a protein null allele (*ss*^*protein null*^) that did not perform transvection (**Fig. S15A-B**). In this background, *Transgene E* expressed Ss in 52% of R7s (**Fig. S15A-B**). We next tested *Transgene E* for transvection with a high-frequency protein null allele (*ss*^*high freq null*^), which can perform transvection to increase *ss* expression (Johnston and Desplan, 2014). When the endogenous *ss* locus was hemizygous for the *ss*^*high freq null*^, we observed no increase in *Transgene E* expression, indicating that it did not perform transvection (51% Ss (Rh4); **Fig. S15A, C**). Moreover, *Transgene E* did not perform transvection in other genetic conditions (**Fig. S15D-E**). Thus, *Transgene E* paired with the endogenous *ss* locus but failed to perform transvection. These data show that an insulator is required for transvection but not for pairing, indicating that *ss* transvection and pairing are mechanistically separable.

### ss pairing and transvection are cell-type-specific

It is poorly understood how pairing impacts transvection in a cell-type-specific manner. We propose two models: constitutive and cell-type-specific buttoning. In the constitutive model, all buttons drive pairing in all cell types, and differences in transvection would occur due to variation in transcription factor binding or chromatin state between cell types (**Fig. 6A**). In the cell-type-specific model, different buttons drive pairing in each cell type, bringing different regions into physical proximity to control transvection efficiency (**Fig. 6A**).

**Figure 6:**
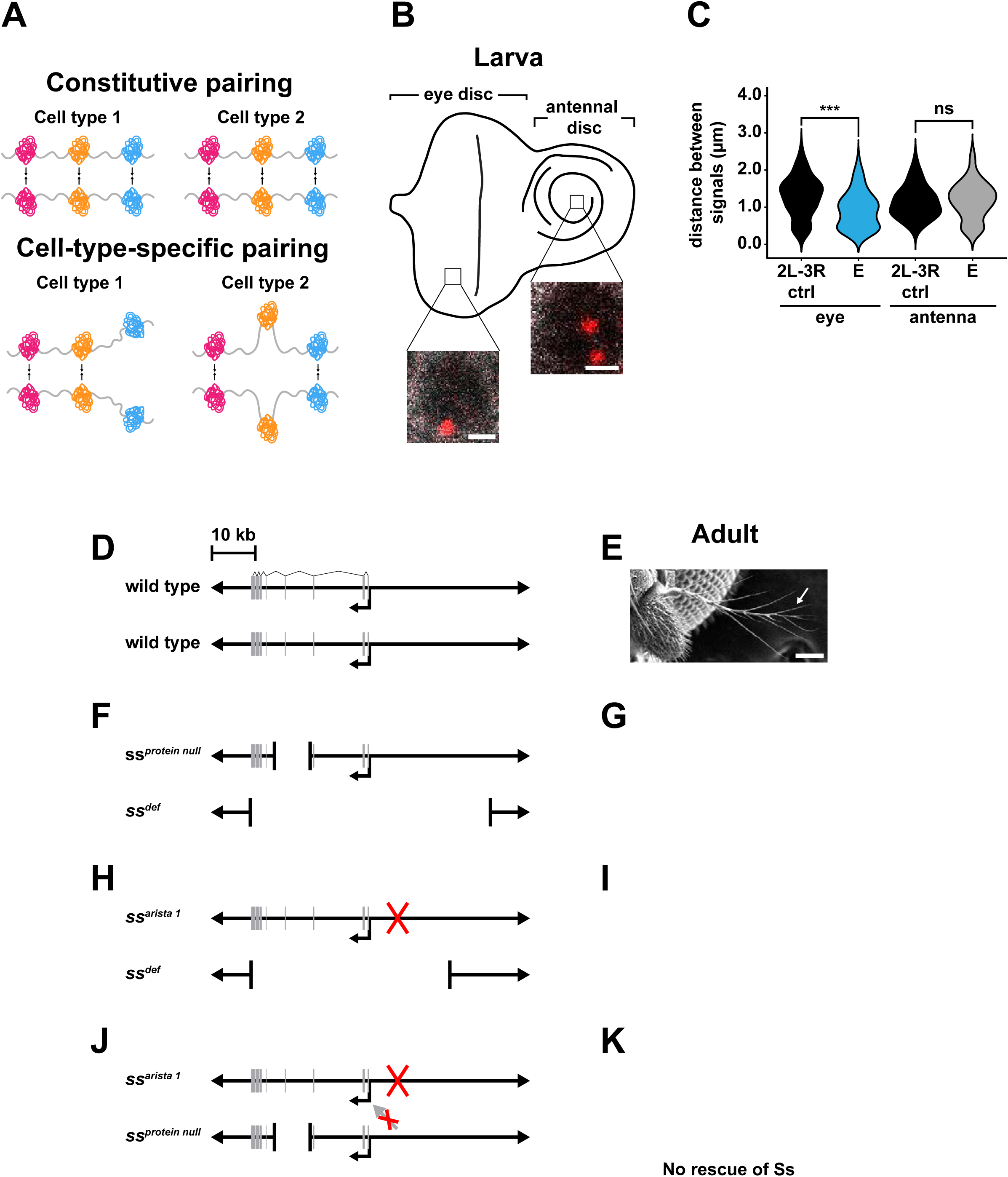
*ss* pairing and transvection are cell-type-specific. **A**. Constitutive vs. cell-type-specific pairing models. **B**. Third instar larval eye-antennal disc. The *ss* button drove pairing in the larval eye but not the larvalantenna. Scale bars=1 μm. White: Lamin B, red: probes against endogenous *ss* and *Transgene E*. **C**. Quantifications for **Fig. 6B**. Black: control, blue: pairer, gray: non-pairer. ***=p<0.05, Wilcoxonrank-sum test; ns=p>0.05, unpaired t-test with Welch’s correction. Data for 2L-3R eye control are thesame as in **Fig. 2F, 3E** (2L-3R control), **3I, S1A-B, S12J**, and **S14Q**. Data for 2L-3R antenna controlare the same as in **Fig. S1A** (2L-3R antenna control). Controls were imaged in two colors, thenpseudocolored red and scored in one color. **D, F, H, J**. Genotypes tested for transvection. Gray rectangles: exons. Smaller black arrows:transcription start sites. Red X indicates an uncharacterized mutation in the *ss*^*arista 1*^ sequence. Red Xover gray arrow indicates an absence of transvection between alleles in the arista. **E, G, I, K**. Arista phenotype. Scale bars=50 μm. White arrows indicate arista. Image for **Fig. 6I** is from the same experiment as **Fig. S17C**.

We tested these models by investigating pairing and transvection of *ss* in two different tissues. In addition to its role in R7 photoreceptors, *ss* is required for the development of the arista, a structure on the antenna (**Fig. 6D-E**) (Morata and Lawrence, 1979; Wernet et al., 2006). *Transgene E*, which contains the *ss* button, drove pairing in the eye but not the antenna from two different insertion sites (sites 1 and 3; **Fig. 6B-C; Fig. S4A-D; Fig. S16A-F**), suggesting that button pairing is cell-type-specific.

As pairing is required for transvection and the *ss* button pairs in a cell-type-specific manner, we hypothesized that transvection at the *ss* locus is cell-type-specific. To test this hypothesis, we examined an allele of *ss* that specifically affects arista development (*ss*^*arista 1*^) (**Fig. 6H; Fig. S17A-F**). In flies transheterozygous for *ss*^*arista 1*^ and a *ss* deficiency (*ss*^*def*^), aristae were transformed into legs (i.e. aristapedia) (**Fig. 6H-I; Fig. S17A, C**). Aristapedia was also observed for *ss*^*protein null*^ / *ss*^*def*^ flies (**Fig. 6F-G**). In the eye, *ss*^*protein null*^ performed transvection to rescue *ss* expression (**Fig. S18A-D**). However, the aristapedia mutant phenotype persisted in *ss*^*arista 1*^ / *ss*^*protein null*^ flies (**Fig. 6J-K; Fig. S17D, F**), suggesting that, unlike in the eye, transvection does not rescue *ss* expression in the arista. Cell-type-specific transvection of the *ss* gene in the eye but not the arista was also observed in other genetic conditions (**Fig. S17G-L; S19A-L**).

As *ss* button pairing and transvection are cell-type-specific and pairing is required for transvection, our data support the cell-type-specific model, in which local buttoning and unbuttoning occur in a cell-type-specific manner to determine transvection efficiency (**Fig. 6A**).

## Discussion

Despite the discovery of homologous chromosome pairing in flies over 100 years ago (Stevens, 1906), the mechanisms that facilitate pairing have remained unclear. We identified multiple button loci interspersed across the genome that drive pairing with their homologous sequences. Specific TADs are responsible for button activity and can pair from multiple locations in the genome. Consistent with our findings, homologous TADs have been observed to interact between paired chromosomes using super resolution microscopy (Szabo et al., 2018). Additionally, homologous chromosome pairing initiates at the same embryonic stage as TAD formation (Dernburg et al., 1996; Fung et al., 1998; Gemkow et al., 1998; Hiraoka et al., 1993; Hug et al., 2017), consistent with a model in which TADs drive homologous chromosomes together.

Our data suggest that complete TADs are sufficient to drive pairing, and that in general, smaller DNA elements such as single insulators do not have strong pairing activity. We propose a model in which complex networks of insulator elements work within the context of a TAD, allowing the domain to take on unique chromatin conformations or bind specific combinations of insulator proteins to create nuclear microcompartments that enable homologous TAD association and pairing.

While TADs are strongly associated with pairing (**Fig. 3F**), a small subset of pairers do not span a TAD (*Transgenes A* and *O*; **Fig. 2F; Fig. S2A; Fig. S5A**), raising the possibility that additional mechanisms work in tandem with TADs to drive homologous chromosomes together(AlHaj Abed, 2018; Erceg, 2018). Additionally, certain transgenes that span TADs do not pair (*Transgenes P* and *BB*; **Fig. 2F; Fig. 3E; Fig. S2B-C; Fig. S5A; Fig. S7C**). These transgenes might span TADs that drive cell-type-specific pairing in non-retinal cell types, similar to *Transgene E*, which drives pairing in photoreceptors but not the antennal disc (**Fig. 6B-C; Fig. S4A-D; Fig. S16A-F**). Alternatively, a subset of TADs may not drive pairing in any cell type.

Our data indicate that pairing and transvection are mechanistically separable: TADs facilitate pairing, while an insulator element facilitates transvection to the endogenous *spineless* locus. Consistent with our findings using endogenous alleles, an insulator is required for transvection but not pairing between transgenes containing the *snail* enhancer and the *eve* promoter (Lim et al., 2018).

We find that the *ss* locus drives pairing and performs transvection in the eye but not in the antenna. Our results support a model in which different buttons drive pairing in different cell types. In this model, local buttoning or unbuttoning at a specific gene determines its transvection efficiency in a given cell type. Variation in levels of pairing or transvection across cell types has been observed for a number of loci (Blick et al., 2016; Fung et al., 1998; Gemkow et al., 1998), suggesting that differences in pairing between cell types may be a general mechanism regulating gene expression.

The mechanisms driving chromosome pairing and transvection have remained a mystery of fly genetics since their initial discoveries by Nettie Stevens and Ed Lewis (Lewis, 1954; Stevens, 1906). Our results provide strong support for the button model of pairing initiation and offer the first evidence of a general feature, specialized TADs, that drives homologous chromosomes together. Furthermore, we find that pairing is necessary but not sufficient for transvection and that distinct elements are required for these processes. Both pairing and transvection are cell-type-specific, suggesting that tighter pairing in a given cell type enables more efficient transvection in that cell type. Our findings suggest a general mechanism in which TADs drive homologous chromosome pairing and interchromosomal gene regulation across organisms to facilitate processes including X-inactivation and imprinting.

**Materials and methods can be found in the supplementary materials**.

## Supplemental Material

### Materials and Methods

#### Drosophila lines

Flies were raised on standard cornmeal-molasses-agar medium and grown at 25° C.

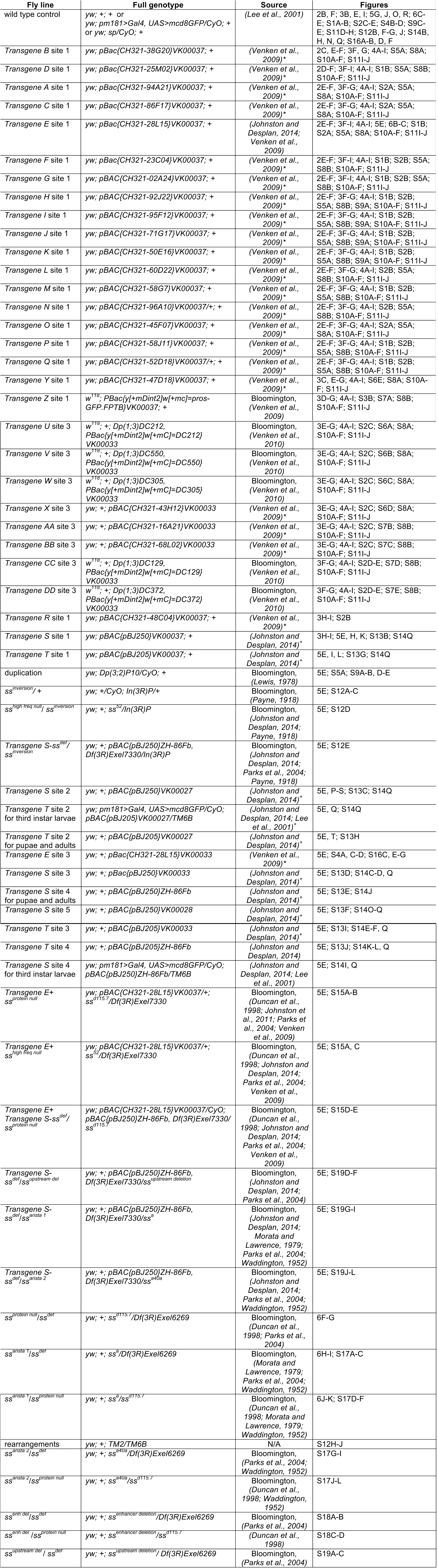

^*^ Constructs were purchased from the CHORI *Drosophila melanogaster* BAC library collection (Venken et al., 2009) and sent to BestGene Inc. (Chino Hills, CA) or Rainbow Transgenic Flies, Inc. (Camarillo, CA) for injection.

^+^ Constructs were generated in (Johnston and Desplan, 2014) and sent to BestGene Inc. (Chino Hills, CA) or Rainbow Transgenic Flies, Inc. (Camarillo, CA) for injection.

Constructs were inserted via PhiC31 integration at the following landing sites:

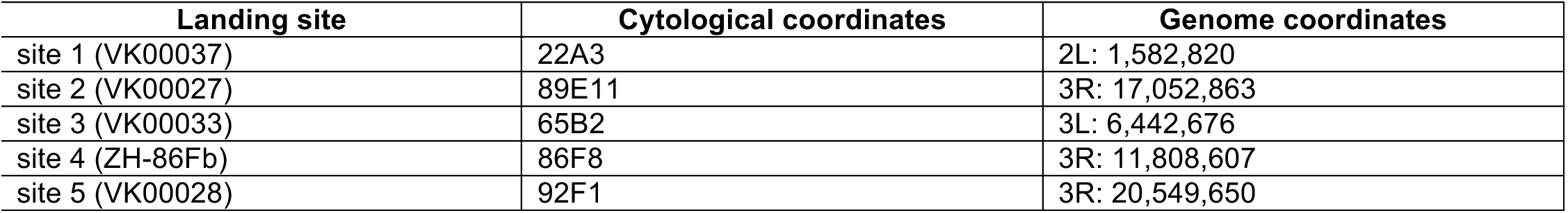

#### Oligopaints probe libraries

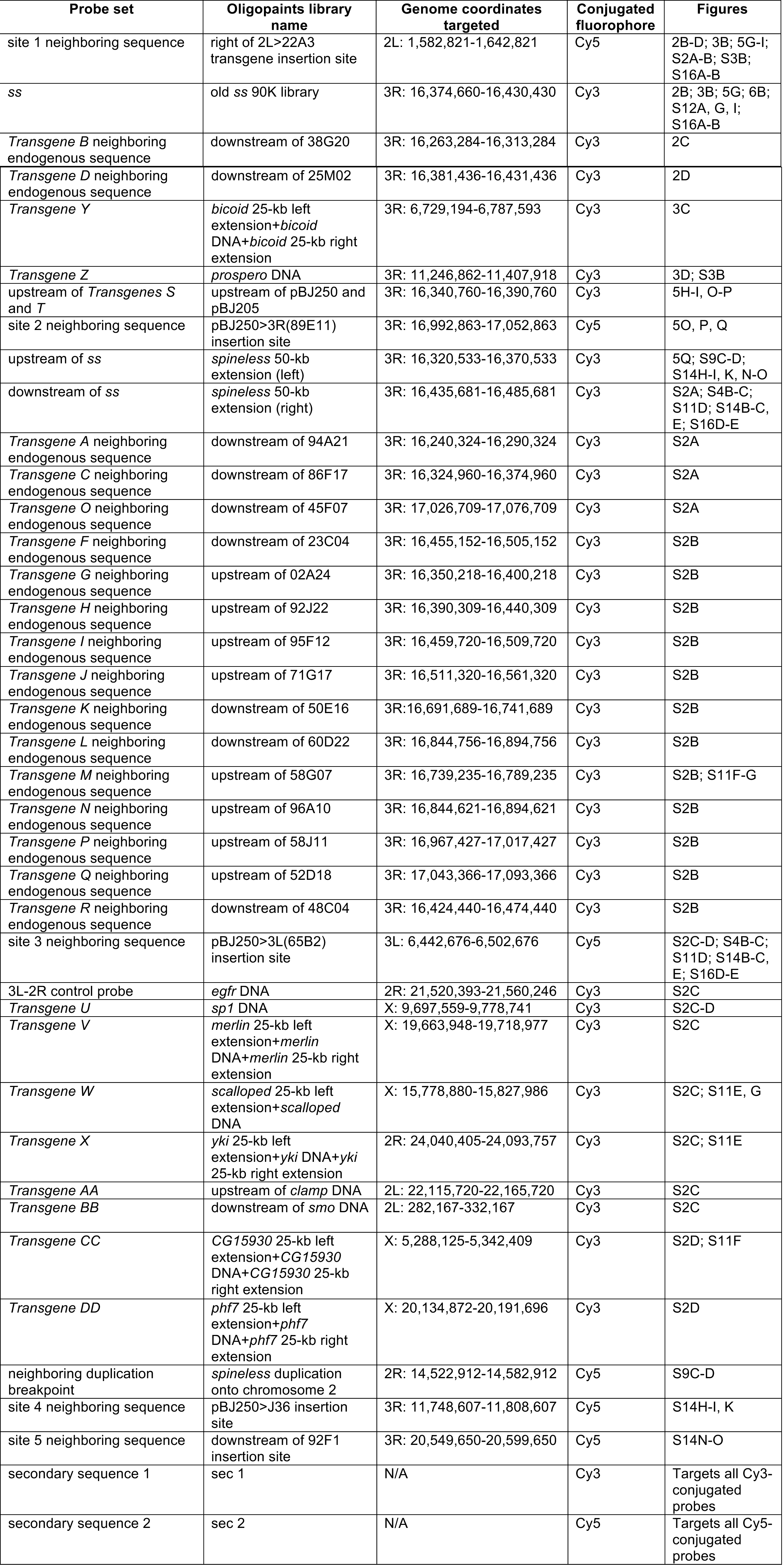

#### Pairing controls

PRs: photoreceptors

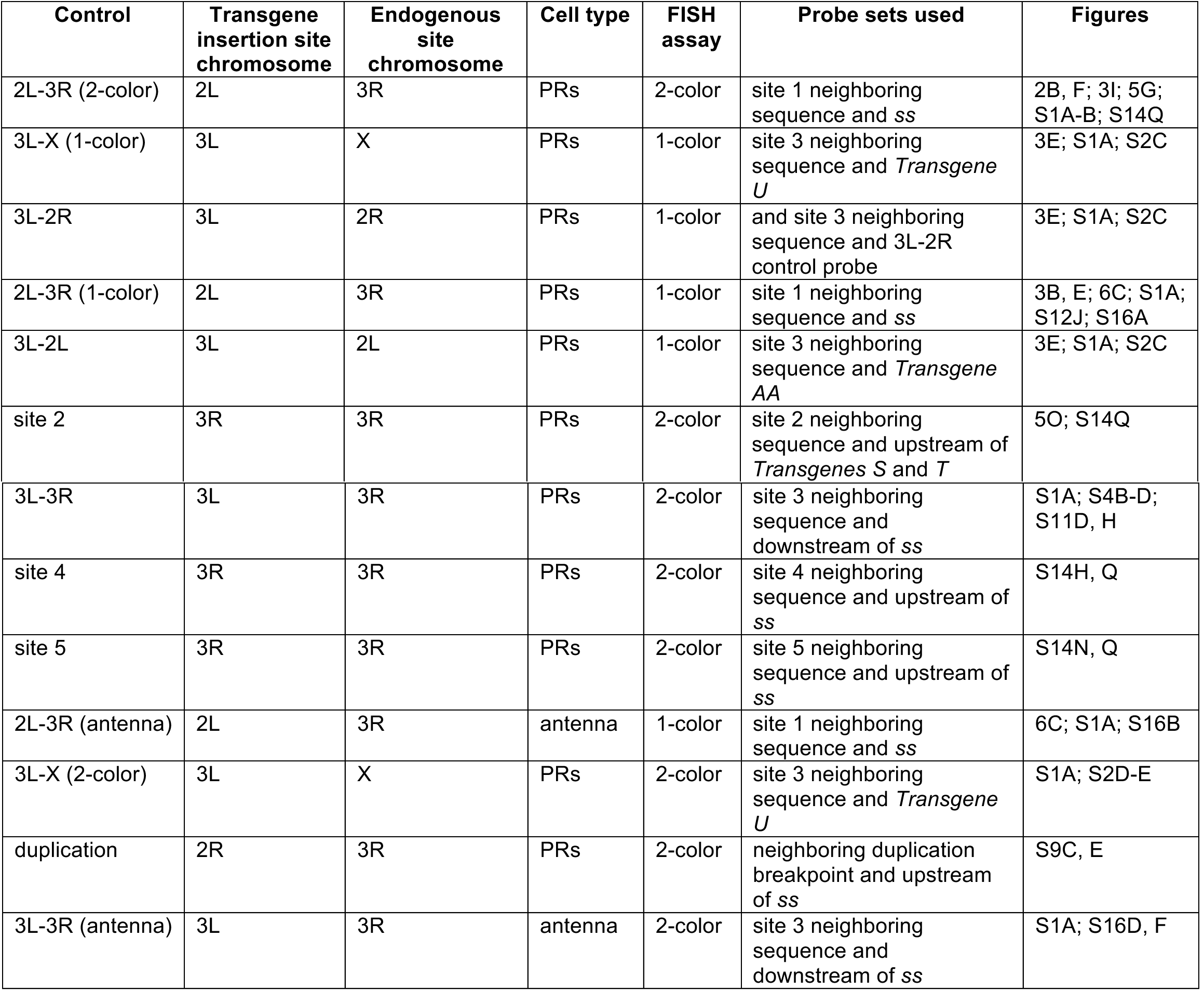

#### Compartments

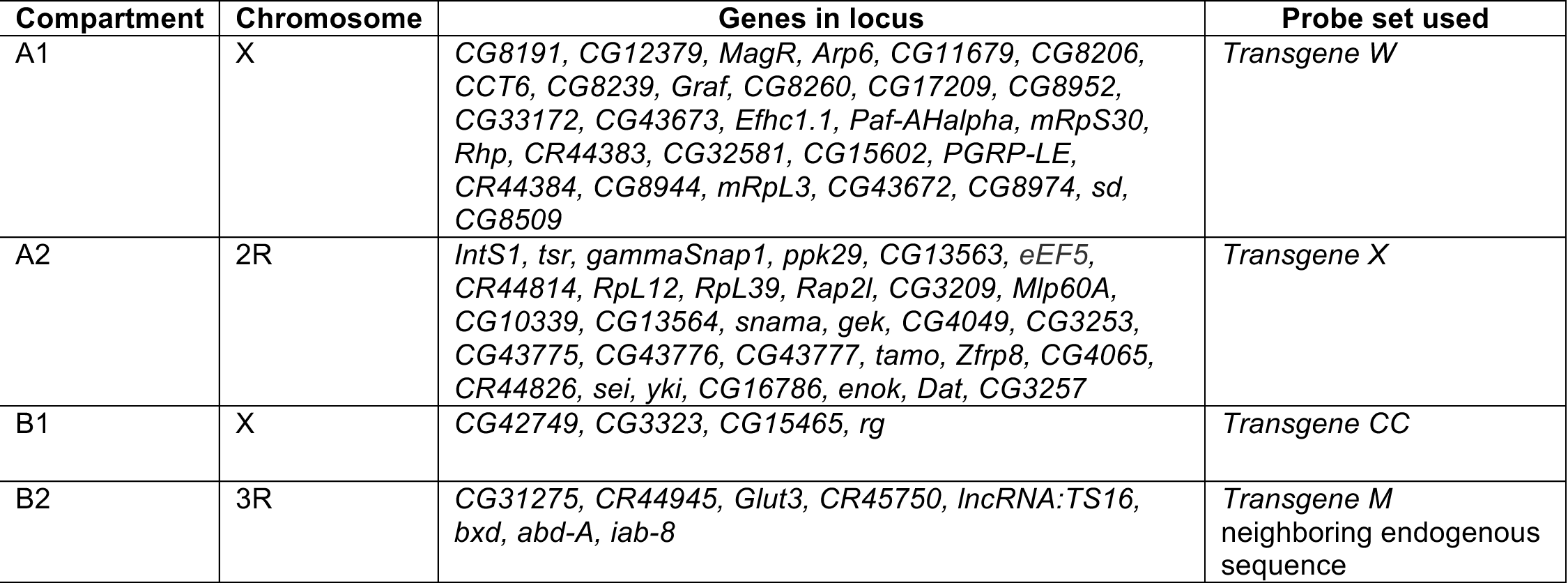

#### Antibodies

Antibodies and dilutions were as follows: mouse anti-Lamin B (DSHB ADL67.10 and ADL84.12), 1:100; rabbit anti-GFP (Invitrogen), 1:500; rabbit anti-Rh4 (gift from C. Zuker, Columbia University), 1:50; mouse anti-Rh3 (gift from S. Britt, University of Texas at Austin), 1:50; mouse anti-Prospero (DSHB MR1A), 1:10; rat anti-Elav (DSHB 7E8A10), 1:50; guinea pig anti-Ss (gift from Y.N. Jan, University of California, San Francisco), 1:500. All secondary antibodies (Molecular Probes) were Alexa Fluor-conjugated and used at a dilution of 1:400.

#### Antibody staining (pupal and adult eyes)

Dissections were performed as described in references (Hsiao et al., 2012; Johnston et al., 2011; Jukam et al., 2016; Thanawala et al., 2013). Eyes were dissected and fixed at room temperature for 15 minutes in 4% formaldehyde diluted in 1X PBX (PBS+0.3% Triton-X), then washed three times in 1X PBX. Eyes were incubated overnight at room temperature in primary antibody diluted in 1X PBX, then washed three times in 1X PBX and incubated in PBX at room temperature for ≥3 hours. Secondary antibody diluted in 1X PBX was added and incubated overnight at room temperature. Eyes were then washed three times in 1X PBX and incubated in PBX at room temperature for ≥2 hours. Adult eyes were mounted in SlowFade Gold (Invitrogen), and pupal eyes were mounted in Vectashield (Vector Laboratories, Inc.). Images were acquired on a Zeiss LSM700 confocal microscope.

The adult eye dissection protocol was used for **Fig. 5D, J-L, R-T; Fig. S12C-E; Fig. S14D, F, J, L, P; Fig. S15B-C, E; Fig. S17B, E, H, K; Fig. S18B, D;** and **Fig. S19B, E, H, K**. The pupal dissection protocol was used for **Fig. 5C** and **Fig. S13B-J**.

#### Oligopaints probe design

Probes for DNA FISH were designed using the Oligopaints technique (Beliveau et al., 2015; Beliveau et al., 2012). Target sequences were run through the bioinformatics pipeline available at http://genetics.med.harvard.edu/oligopaints/ to identify sets of 42-bp (for old ss 90K probes) or 50-bp (for all other probes) optimized probe sequences (i.e. “libraries”) tiled across the DNA sequence of interest. Five 19-bp barcoding primers, gene F and R; universal (univ) F and R, and either sublibrary (sub) F or random (rando) R, were appended to the 5’ and 3’ ends of each probe sequence (**Fig. S20A-B**). To ensure that all probes were the same length, an additional 8-bp random sequence was added to the 3’ end of the old ss 90K probes. The gene F and R primers allowed PCR amplification of a probe library of interest out of the total oligo pool, and the univ F and R primers allowed conjugation of fluorophores, generation of single-stranded DNA probes, and PCR addition of secondary sequences to amplify probe signal. The *ss* 50-kb left and right extension libraries had a sub F primer between the gene and universal forward primers to allow PCR amplification of probes targeting a specific sub-region of the locus of interest (**Fig. S20A**). All other probe libraries had a rando R primer appended at the 3’ end to maintain a constant sequence length between all probes (**Fig. S20B**).

Barcoding primer sequences were taken from a set of 240,000 randomly generated, orthogonal 25-bp sequences (Qikai Xu, 2008) and run through a custom script to select 19-bp sequences with ≤15-bp homology to the *Drosophila* genome. Primers were appended to probe sequences using the orderFile.py script available at http://genetics.med.harvard.edu/oligopaints/. Completed probe libraries were synthesized as custom oligo pools by Custom Array, Inc. (Bothell, WA), and fluorescent FISH probes were generated as described in references (Beliveau et al., 2015; Beliveau et al., 2012).

#### DNA FISH

DNA FISH was performed using modified versions of the protocols described in references (Beliveau et al., 2015; Beliveau et al., 2012). 20-50 eye-antennal discs attached to mouth hooks from third instar larvae were collected on ice and fixed in 129 μL ultrapure water, 20 μL 10X PBS, 1 μL Tergitol NP-40, 600 μL heptane, and 50 μL fresh 16% formaldehyde. Tubes containing the fixative and eye discs were shaken vigorously by hand, then fixed for 10 minutes at room temperature with nutation. Eye discs were then given three quick washes in 1X PBX, followed by three five-minute washes in PBX at room temperature with nutation. Eye discs were then removed from the mouth hooks and blocked for 1 hour in 1X PBX+1% BSA at room temperature with nutation. They were then incubated in primary antibody diluted in 1X PBX overnight at 4°C with nutation. Next, eye discs were washed three times in 1X PBX for 20 minutes and incubated in secondary antibody diluted in 1X PBX for two hours at room temperature with nutation. Eye discs were then washed two times for 20 minutes in 1X PBX, followed by a 20-minute wash in 1X PBS. Next, discs were given one 10-minute wash in 20% formamide+2X SSCT (2X SSC+.001% Tween-20), one 10-minute wash in 40% formamide+2X SSCT, and two 10-minute washes in 50% formamide+2X SSCT. Discs were then predenatured by incubating for four hours at 37°C, three minutes at 92°C, and 20 minutes at 60°C. Primary probes were added in 45 μL hybridization buffer consisting of 50% formamide+2X SSCT+2% dextran sulfate (w/v), + 1 μL RNAse A. All probes were added at a concentration of ≥5 pmol fluorophore/μL. For FISH experiments in which a single probe was used, 4 μL of probe was added. For FISH experiments in which two probes were used, 2 μL of each probe was added. After addition of probes, eye discs were incubated at 91°C for three minutes and at 37°C for 16-20 hours with shaking. Eye discs were then washed for 1 hour at 37°C with shaking in 50% formamide+2X SSCT. 1 μL of each secondary probe was added at a concentration of 100 pmol/μL in 50 μL of 50% formamide+2X SSCT. Secondary probes were hybridized for 1 hour at 37°C with shaking. Eye discs were then washed twice for 30 minutes in 50% formamide+2X SSCT at 37°C with shaking, followed by three 10-minute washes at room temperature in 20% formamide+2X SSCT, 2X SSCT, and 2X SSC with nutation. Discs were mounted in SlowFade Gold immediately after the final 2X SSC wash, and imaged using a Zeiss LSM700 confocal microscope.

#### Generation of CRISPR lines

CRISPR lines were generated as described in references (Anderson et al., 2017; Gratz et al., 2013; Port et al., 2014; Yan et al., 2017). For both *ss*^*enh del*^ and *ss*^*upstream del*^, sense and antisense DNA oligos for the forward and reverse strands of four gRNAs were designed to generate BbsI restriction site overhangs. The oligos were annealed and cloned into the pCFD3 cloning vector (Addgene, Cambridge, MA). A single-stranded DNA homology bridge was generated with 60-bp homologous regions flanking each side of the predicted cleavage site and an EcoRI (for *ss*^*enh del*^) or NaeI (for *ss*^*upstream del*^) restriction site to aid in genotyping. The gRNA constructs (125 ng/μl) and homologous bridge oligo (100 ng/μl) were injected into *Drosophila* embryos (BestGene, Inc., Chino Hills, CA). Single males were crossed with a balancer stock (*yw; +; TM2/TM6B*), and F1 female progeny were screened for the insertion via PCR, restriction digest, and sequencing. Single F1 males whose siblings were positive for the deletion were crossed to the balancer stock (*yw; +; TM2/TM6B*), and the F2 progeny were screened for the deletion via PCR, restriction digest, and sequencing. Deletion-positive flies from multiple founders were used to establish independent stable stocks.

The following oligos were used for the *ss*^*enh del*^ CRISPR:

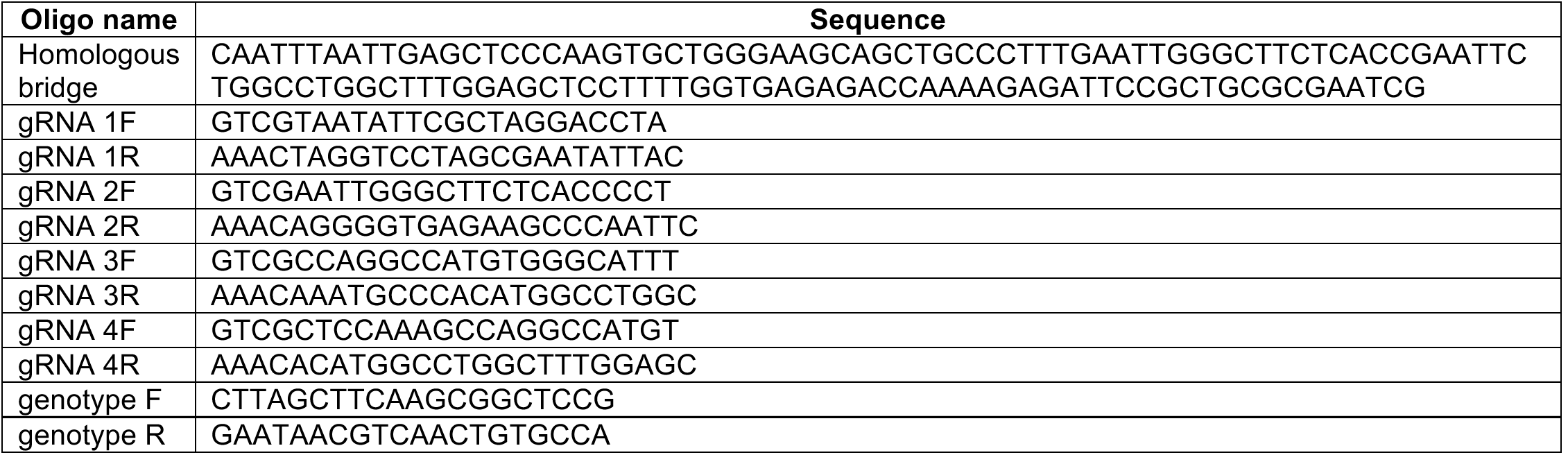

The following oligos were used for the *ss*^*upstream del*^ CRISPR:

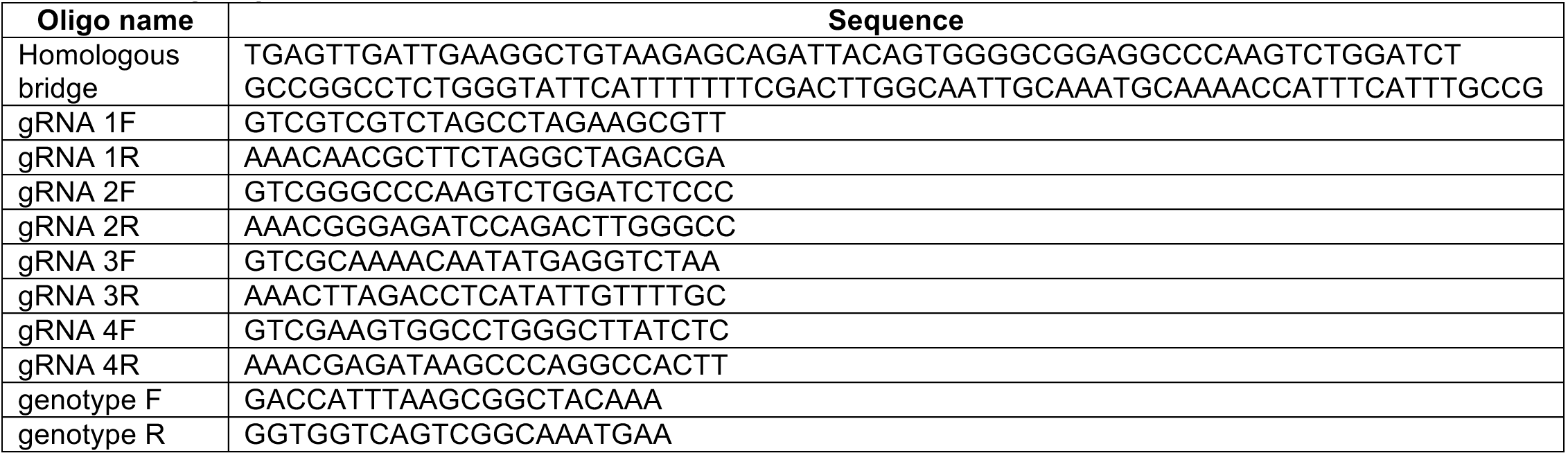

#### Scanning electron microscopy

Adult *Drosophila* heads were removed and immediately mounted on a pin stub without fixation or sputtering. Heads were imaged at high vacuum at a voltage of 1.5 kV. All SEM was performed on a FEI Quanta ESEM 200 scanning electron microscope. SEM was used for **Fig. 6E, G, I, K; Fig. S17C, F, I, L;** and **Fig. S19C, F, I, L**.

#### Pairing quantifications

All quantifications were performed in 3D on z-stacks with a slice thickness of 0.2 μm. Quantifications were performed manually using Fiji (Schindelin et al., 2012; Schneider et al., 2012). To chart the z position of each FISH dot, a line was drawn through the dot and the Plot Profile tool was used to assess the stack in which the dot was brightest. To determine the x-y distance between the two FISH dots, a line was drawn from the center of one dot to the center of the other dot and the length of the line was measured with the Plot Profile tool. The distance between the FISH dots was then calculated in 3D. A total of 50 nuclei from three eye discs were quantified for each genotype (i.e. N=3, n=50).

For experiments in which the transgene and endogenous site were both labeled with red fluorescent probes, FISH punctae ≤0.4 μm apart could not be distinguished as separate and were assigned a distance of 0.4 μm apart. For all controls in **Fig. 3E**, **6C**, and **S12J**, green probes labeling the transgene insertion site were pseudocolored red and data were quantified in the same way as experiments in which the transgene and endogenous site were both labeled with red probes. 3L-X control data in **Fig. 3E** are taken from the same experiment as in **Fig. S2E**, but the data were requantified with the green probes pseudocolored red. Similarly, 2L-3R eye control data in **Fig. 3E, 6C**, and **S12J** are taken from the same experiment as in **Fig. 2F, 3I, S1A-B**, and **S14Q**, but the data were re-quantified with the green probes pseudocolored red.

#### Adult eye quantifications

The frequencies of Rh4-and Rh3-expressing R7s were scored manually for at least eight eyes per genotype. R7s co-expressing Rh3 and Rh4 were scored as Rh4-positive. 100 or more R7s were scored for each eye. For **Fig. S19E, H**, and **K**, only the ventral half of each eye was scored.

#### Hi-C mapping and TAD calling

Directionality index scores were calculated across 15-kb windows, stepping every 5 kb, by finding the log2 transform of the difference in the ratios of downstream versus upstream summed observed over expected interactions ranging from 15 kb to 100 kb in size. The expected value of a bin was defined as the sum of the product of fragment corrections for each valid fragment pair with both interaction fragments falling within the bin.

Directionality indices were generated using 14 published Hi-C datasets (Hou et al., 2012; Li et al., 2015; Schuettengruber et al., 2014; Stadler et al., 2017):

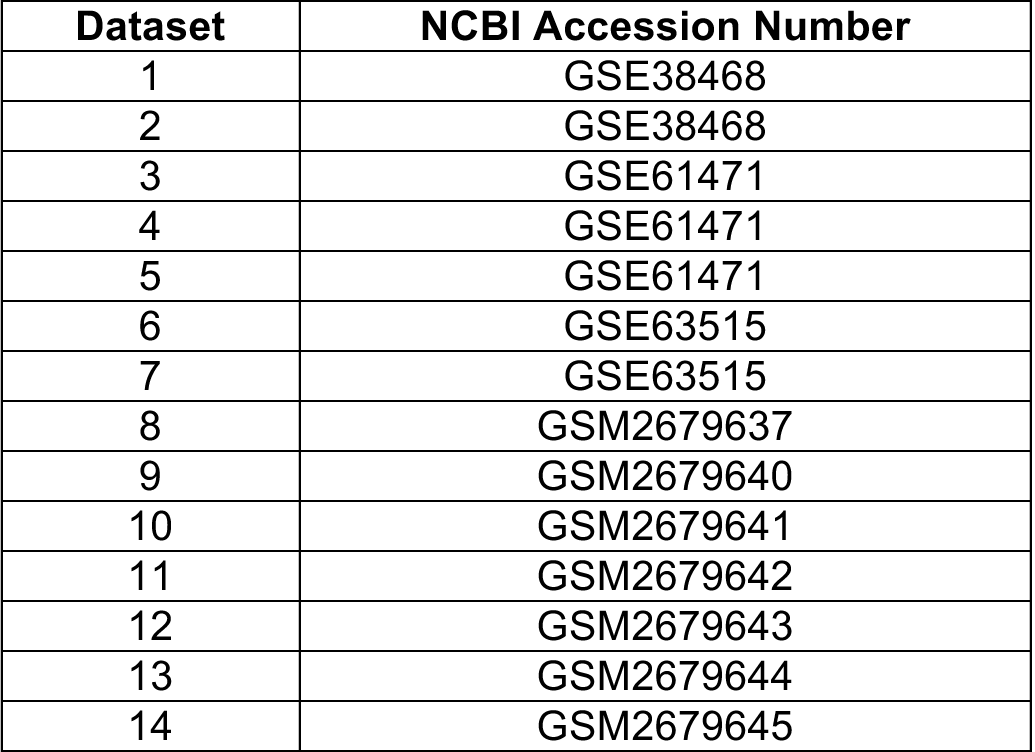

TADs were read from the beginning of a positive directionality index peak to the end of a negative directionality index peak. Parameters for calling a TAD were as follows: **1)** The positive peak must have a signal of ≥0.8; **2)** The negative peak must have a signal of ≤0.8; and **3)** The TAD must be present in at least two datasets. Any transgene covering ≥95% of a TAD was considered to span a TAD.

#### mRNA sequencing and analysis

RNA-seq was performed on three biological replicates, each consisting of 30 third instar larval eye discs. Eye discs were dissected in 1X PBS, separated from the mouth hooks and antennal discs, and placed directly into 300 μL of Trizol. RNA was purified using a Zymo Direct-zol RNA MicroPrep kit (catalog number R2062). mRNA libraries were prepared using an Illumina TruSeq Stranded mRNA LT Sample Prep Kit (catalog number RS-122-2101). Sequencing was performed using an Illumina NextSeq 500 (75 bp, paired end). Sequencing returned an average of 23,048,349 reads per replicate.

The following pipeline was used for mRNA-sequencing analysis: 1) FASTQ sequencing datasets were assessed for quality using FastQC; 2) Pseudoalignment with the *Drosophila* dm6 transcriptome and read quantifications were performed using kallisto (Bray et al., 2016); 3) Transcript abundance files generated by kallisto were joined to a file containing the genomic coordinates of all *Drosophila* mRNA transcripts (dmel-all-r6.20.gtf, available from Flybase); 4) The joined transcript coordinate file was compared to a file containing the coordinates of all tested transgenes using the bedtools intersect tool (http://bedtools.readthedocs.io/en/latest/content/tools/intersect.html)(Quinlan and Hall, 2010). The output file contained a list of all TPMs for each gene contained in each transgene.

#### Assessment of chromatin marks and ncRNA, Polycomb Group Complex, and insulator density

ncRNA content of transgenes was assessed manually using the GBrowse tool on FlyBase. tRNAs, miRNAs, snoRNAs, and lncRNAs were included in the analysis of ncRNA content.

Transgenes were evaluated for insulator binding sites, Polycomb Group Complex binding sites, and the presence of chromatin marks using publicly available ChIP-chip and ChIP-seq datasets(Cuartero et al., 2014; Negre et al., 2010; Ong et al., 2013; Van Bortle et al., 2012; Wood et al., 2011). The following datasets were used for this analysis:

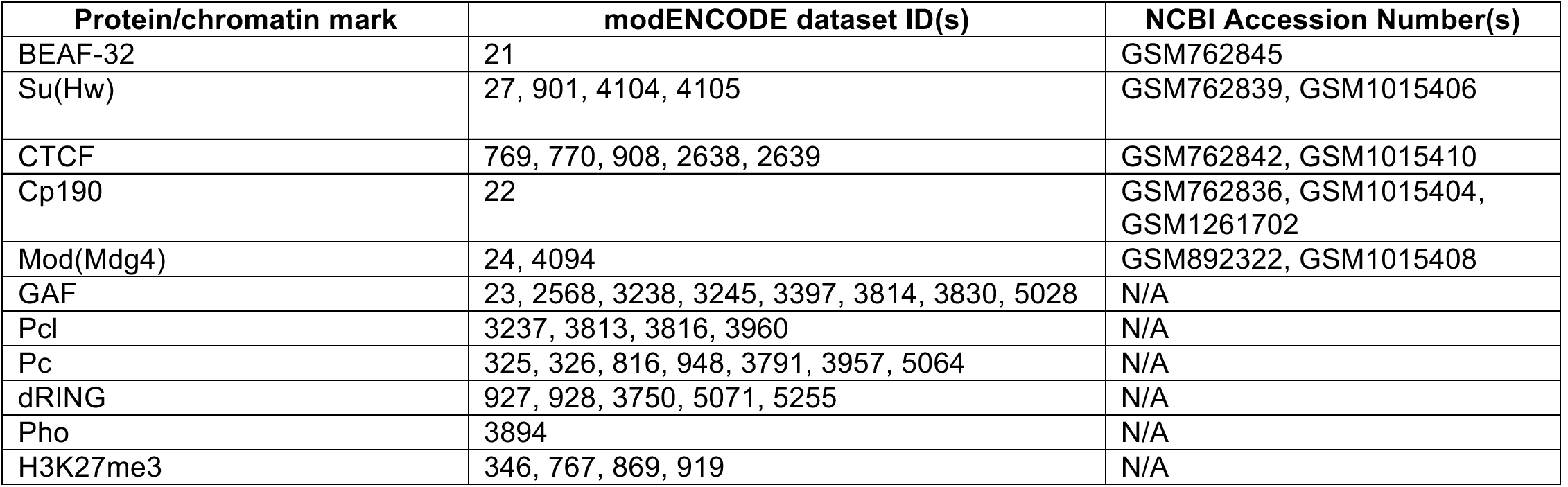

To ensure a higher likelihood of selecting true ChIP peaks rather than false positives, only those insulators present in data from multiple cell types were considered when assessing the number of insulator sites per transgene‥bed files containing the genomic coordinates of all ChIP peaks in each dataset were downloaded and classified by cell type. All datasets from the same cell type were merged into one file using the bedtools merge tool(http://bedtools.readthedocs.io/en/latest/content/tools/merge.html)(Quinlan and Hall, 2010). Insulator ChIP peaks present in more than one cell type were identified using the bedtools multiIntersectBed tool(Quinlan and Hall, 2010). The coordinates of insulator ChIP peaks present in multiple cell types were compared to a .bed file containing the genomic coordinates of all transgenes using the bedtools intersect tool (http://bedtools.readthedocs.io/en/latest/content/tools/intersect.html)(Quinlan and Hall, 2010). This pipeline output the number of insulator ChIP peaks contained in each transgene.

To identify clusters of insulators, files containing the ChIP peak coordinates for each insulator were compared using the the bedtools multiIntersectBed tool(Quinlan and Hall, 2010), which output a .bed file listing all of the locations of overlap between insulator binding sites. Insulators were considered to cluster if their binding site coordinates overlapped or were directly adjacent to each other. The coordinates of clusters containing specific numbers or combinations of insulators were selected from the intersected .bed file and compared to a .bed file containing the genomic coordinates of all transgenes using the bedtools intersect tool(http://bedtools.readthedocs.io/en/latest/content/tools/intersect.html)(Quinlan and Hall, 2010). This pipeline output the number of insulator clusters contained in each transgene.

For Polycomb Group Complex proteins and chromatin marks, .bed files containing the genomic coordinates of all ChIP peaks in each dataset were downloaded and merged into one file using the bedtools merge tool(http://bedtools.readthedocs.io/en/latest/content/tools/merge.html)(Quinlan and Hall, 2010). The merged file was compared to a .bed file containing the genomic coordinates of all transgenes using the bedtools intersect tool (http://bedtools.readthedocs.io/en/latest/content/tools/intersect.html)(Quinlan and Hall, 2010). This pipeline output the number of protein or chromatin mark ChIP peaks contained in each transgene.

#### Statistical analysis

All datasets were tested for a Gaussian distribution using a D’Agostino and Pearson omnibus normality test and a Shapiro-Wilk normality test. If either test indicated a non-Gaussian distribution for any of the datasets in an experiment, datasets were tested for statistical significance using a Wilcoxon rank-sum test (for single comparisons) or a one-way ANOVA on ranks with Dunn’s multiple comparisons test (for multiple comparisons). If both the D’Agostino and Pearson and the Shapiro-Wilk tests indicated a Gaussian distribution for all datasets in an experiment, datasets were tested for statistical significance using an unpaired t-test with Welch’s correction (for single comparisons) or an ordinary one-way ANOVA with Dunnett’s multiple comparisons test (for multiple comparisons).

*Maximum likelihood calculations*: Parameters for either a single or double Gaussian distribution were estimated using maximum likelihood, and model selection was subsequently performed using the Bayesian Information Criterion (BIC). For the double Gaussian, parameters were estimated using a nonlinear recursion (Levenberg-Marquardt) algorithm to maximize the log likelihood of the distribution.

Maximum likelihood calculations were performed for all transgenes tested in **Fig. 2F**. Maximum likelihood estimation was not possible for transgenes in **Fig. 3E**, as the 0.4 μm distance cutoff for 1-color FISH was too high to allow separation of paired and unpaired distributions into two Gaussian distributions.

## Supplemental Figure Legends

**Supplemental Figure 1:**
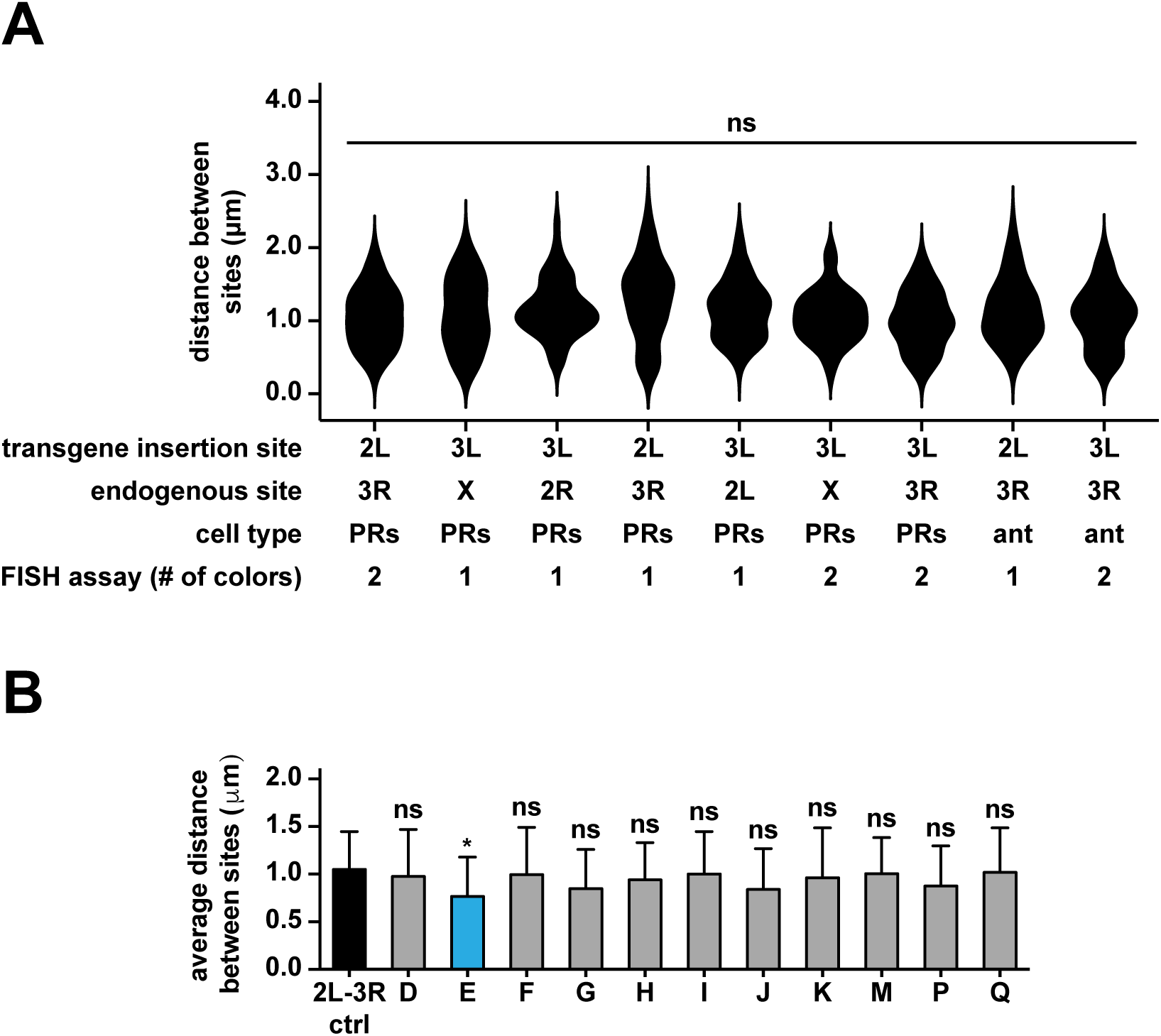
Comparisons between pairing controls and additional statistical tests confirm identification of pairers. **A**. Quantifications for all negative controls used to assess pairing between a transgene and its endogenous site in this study. PRs: photoreceptors, ant: antenna. ns=p>0.05, one-way ANOVA on ranks with Dunn’s multiple comparisons test. Data for 2L-3R PRs 2-color control are the same as in**Fig. 2F, 3I, S1B**, and **S14Q** (2L-3R control). Data for 3L-X 1-color control are the same as in **Fig. 3E**(3L-X control). Data for 3L-2R control are the same as in **Fig. 3E** (3L-2R control). Data for 2L-3R PRs1-color control are the same as in **Fig. 3E** (2L-3R control), **6C** (2L-3R eye control), and **S12J**. Data for3L-2L control are the same as in **Fig. 3E** (3L-2L control). Data for 3L-X 2-color control are the sameas in **Fig. S2E**. Data for 3L-3R control are the same as in **Fig. S4D, S11H, S14Q** (3L-3R control).Data for 2L-3R antenna control are the same as in **Fig. 6C** (2L-3R antenna control). Data for 3L-3Rantenna are the same as in **S16F**. All 1-color controls were imaged in two colors, then pseudocolored red and scored in one color. **B**. Comparison of the means for all datasets in **Fig. 2F** fit to a single Gaussian distribution by maximum likelihood estimation. Black: control, blue: pairers, gray: non-pairers. *=p<0.05, ns=p>0.05,ordinary one-way ANOVA with Dunnett’s multiple comparisons test. Data for 2L-3R control are the same as in **Fig. 2F, 3E** (2L-3R control), **6C** (2L-3R eye control), **S1A, S12J**, and **S14Q** (2L-3Rcontrol). Data for *Transgene D-G* are the same as in **Fig. 2A** and **3I**. Data for *Transgene H-K, M, P*,and *Q* are the same as in **Fig. 2A**.

**Supplemental Figure 2:**
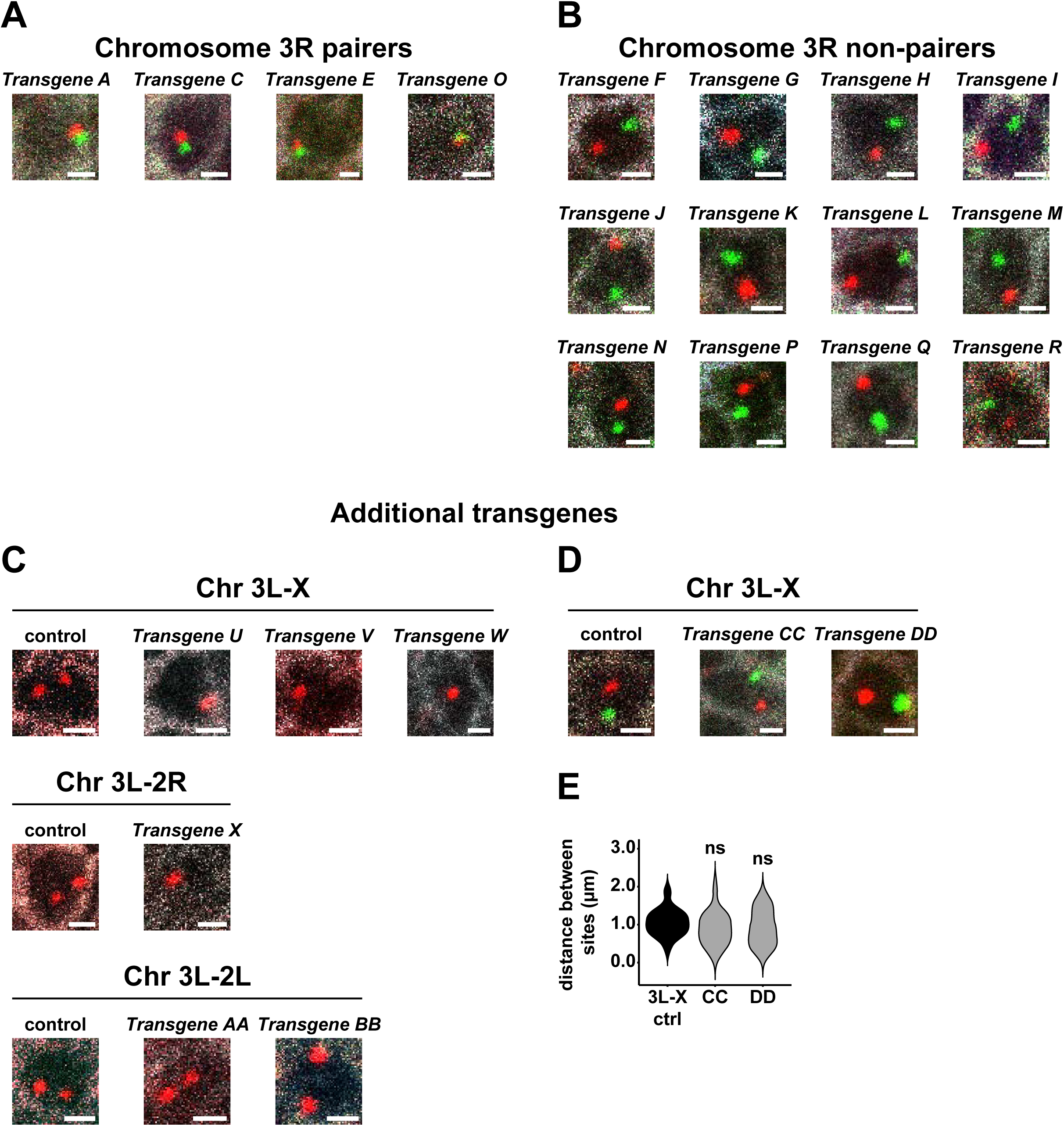
A subset of transgenes interspersed across the genome drive pairing. **A-B**. Pairer and non-pairer transgenes from the initial screen of a ~1 Mb region of chromosome 3R. Scale bars=1 μm. White: Lamin B, red: probes neighboring endogenous sequence, green: probes neighboring transgene insertion site. **C**. Additional transgenes taken from chromosomes X (*Transgenes U-W*), 2R (*Transgene X*), and 2L(*Transgenes AA and BB*) and inserted at site 3 on Chr 3L. 3L-X control image is from the same experiment as **Fig. S2D** control. **D**. Additional transgenes taken from chromosome X and inserted at site 3. Pairing was assessed witha two-color FISH strategy. Scale bars=1 μm. White: Lamin B, red: probes neighboring endogenous sequence, green: probes neighboring transgene insertion site. Control image is from the same experiment as **Fig. S2C** (3L-X control). **E**. Quantifications for transgenes in **Fig. S2D**. Neither transgene contained a TAD. Black: control, gray: non-pairers. ns=p>0.05, one way ANOVA on ranks with Dunn’s multiple comparisons test. Control data are the same as in **Fig. 3E** (3L-X control) and **S1A**.

**Supplemental Figure 3:**
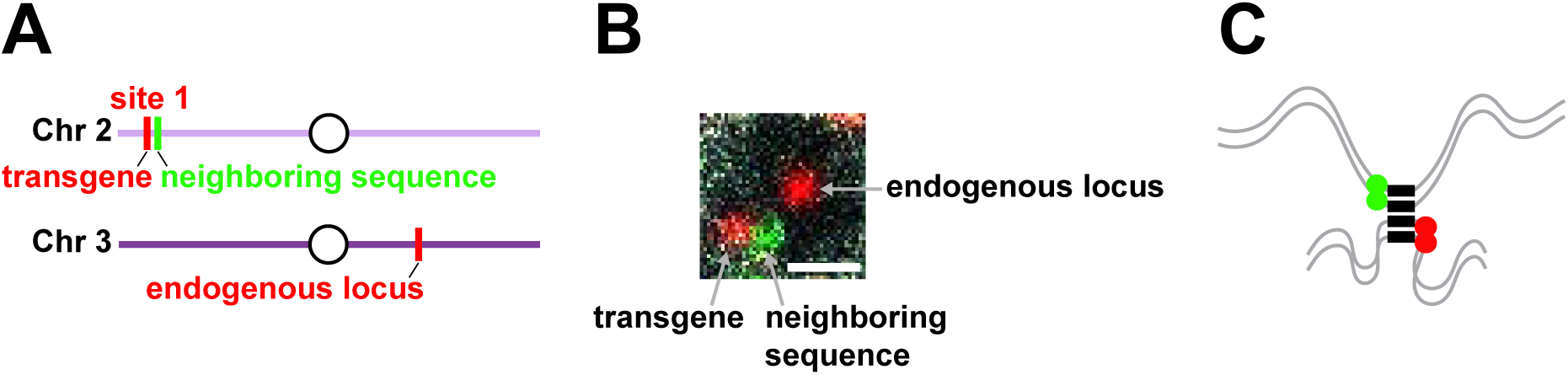
Probes neighboring paired sequences give offset probe signals. **A**. Schematic of FISH strategy used to label transgene and endogenous sequences and the region directly neighboring the transgene insertion site. **B**. Probes directly neighboring the transgene insertion site could be distinguished from probes labeling the transgene sequence itself, despite being immediately downstream on the DNA. Image is from the same experiment as **Fig. 3D**. **C**. When a transgene drives pairing with its endogenous site, the two copies of the transgene are paired with each other and the two copies of the endogenous site are paired with each other due to homologous chromosome pairing. Therefore, one green FISH puncta (neighboring the transgene insertion site) and one red FISH puncta (neighboring the endogenous site) are observed. The experiment in **Fig. S3B** showed that two sets of probes targeting neighboring regions on the DNA could be distinguished from each other. As the red and green probes in **Fig. S3C** are neighboring the paired sites, not directly targeting the paired sites, it is therefore feasible that their signals do not completely overlap, despite being close in 3D space.

**Supplemental Figure 4:**
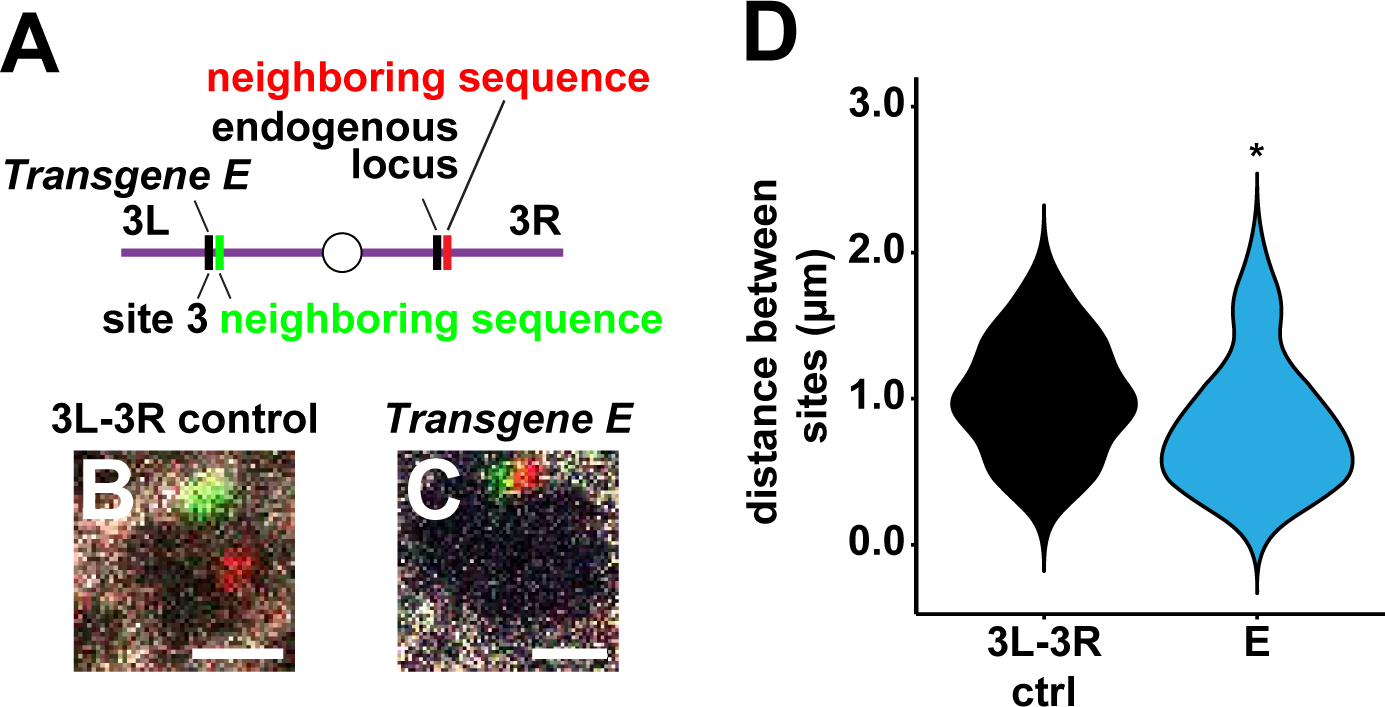
Buttons drive pairing in a position-independent manner. **A**. DNA FISH strategy used to assess pairing of *Transgene E* inserted at site 3 with its endogenous locus. **B-C**. Control and *Transgene E* at site 3. Scale bars=1 μm. White: Lamin B, red: probes neighboring endogenous sequence, green: probes neighboring transgene insertion site. Image for **S4B** is from the same experiment as in **S11D** and **S14B**. **D**. Quantifications for **Fig. S4B-C**. Black: control, blue: pairer. *=p<0.05, Wilcoxon rank-sum test.Data for control are the same as in **Fig. S1A** (3L-3R control), **S11H**, and **S14Q** (3L-3R control).

**Supplemental Figure 5:**
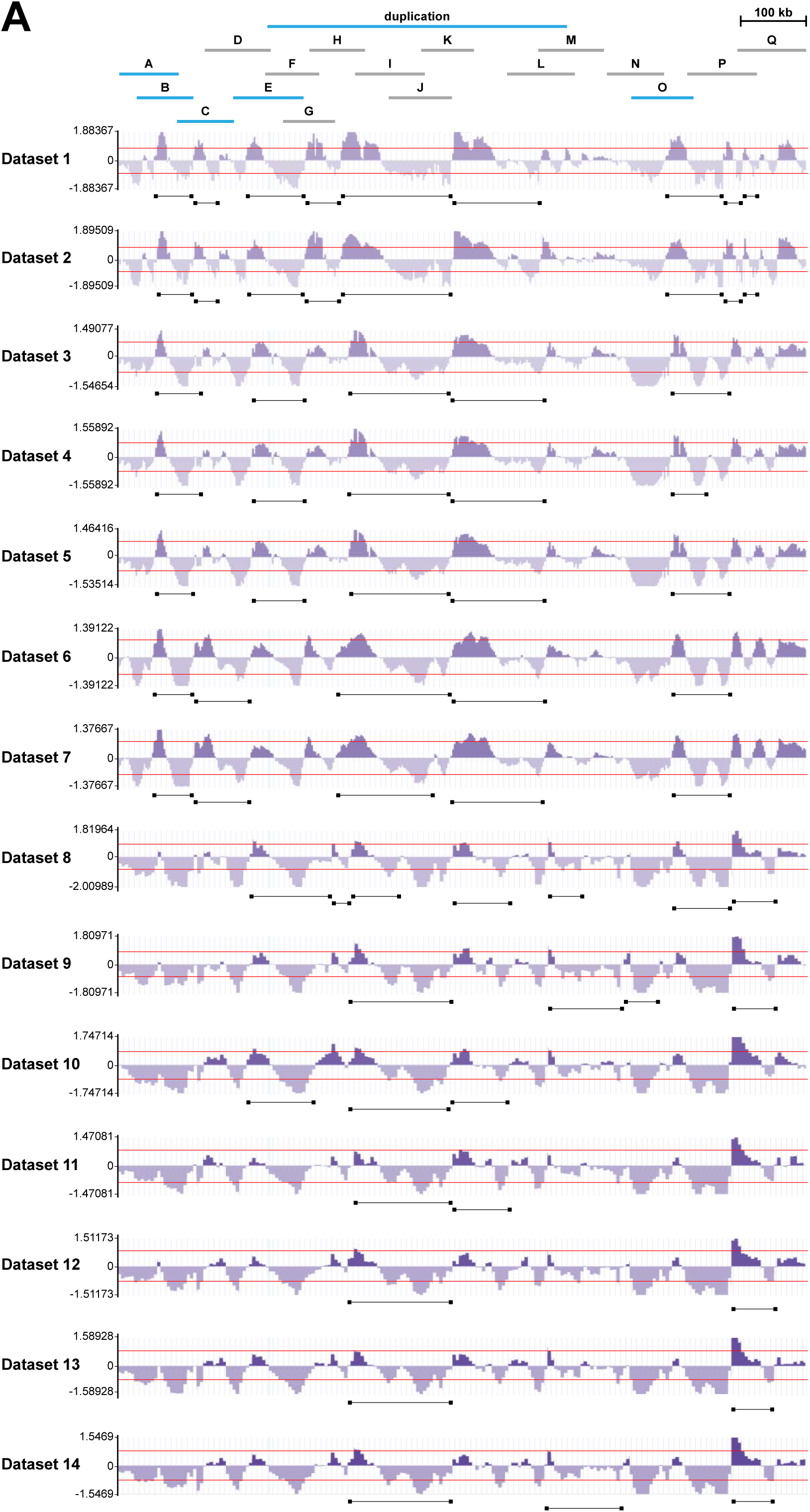
TAD calls across 14 Hi-C datasets for the region on chromosome 3R used for the initial pairing screen. **A**. Red lines indicate a directionality index signal of 0.8 or −0.8, the cutoff for a TAD. Black bars indicate TAD calls.

**Supplemental Figure 6:**
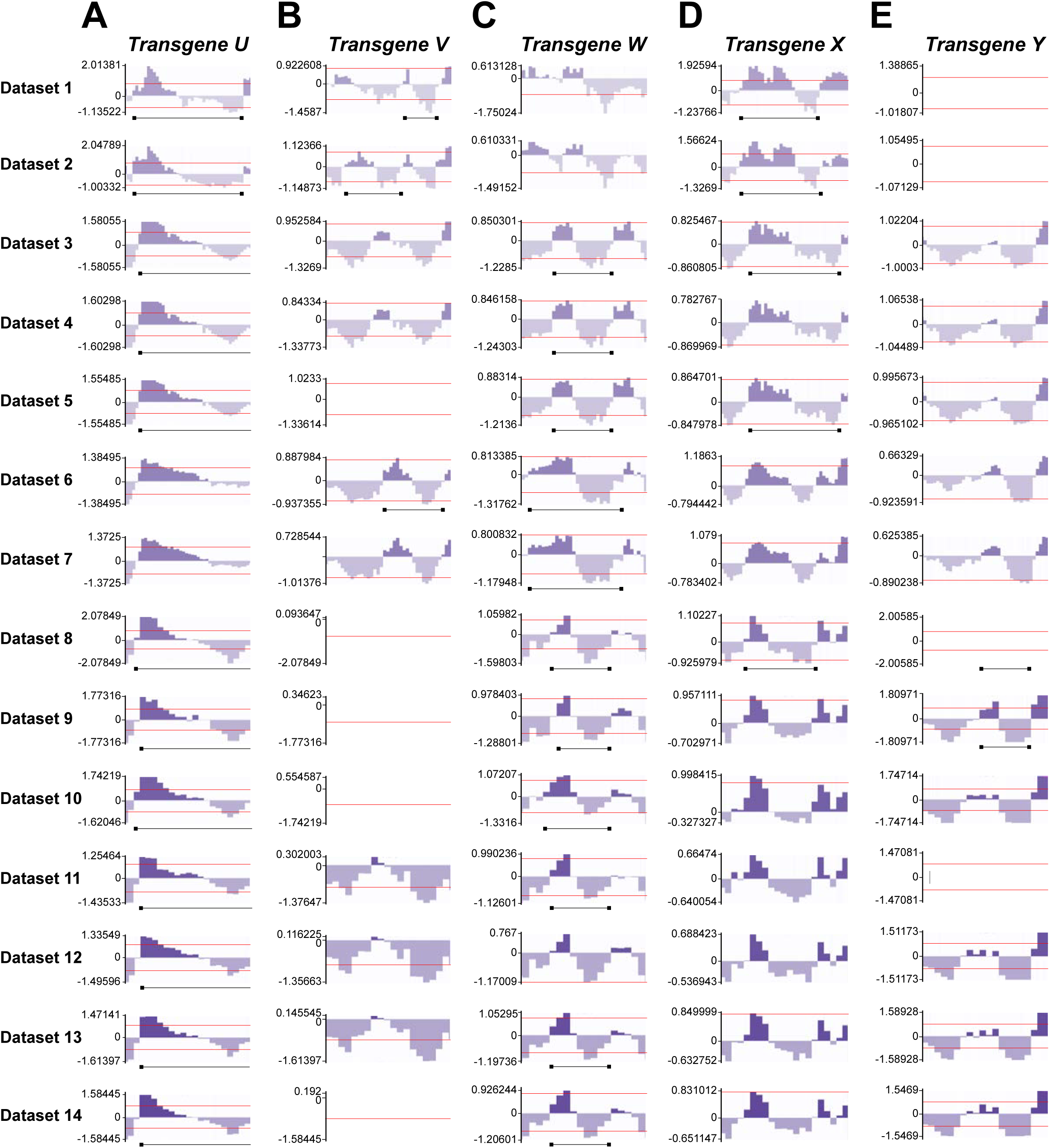
TAD calls across 14 Hi-C datasets for Transgenes U-Y. **A-E**. Red lines indicate a directionality index signal of 0.8 or −0.8, the cutoff for a TAD. Black bars indicate TAD calls. Directionality indices are shown for the entire region spanned by each transgene.

**Supplemental Figure 7:**
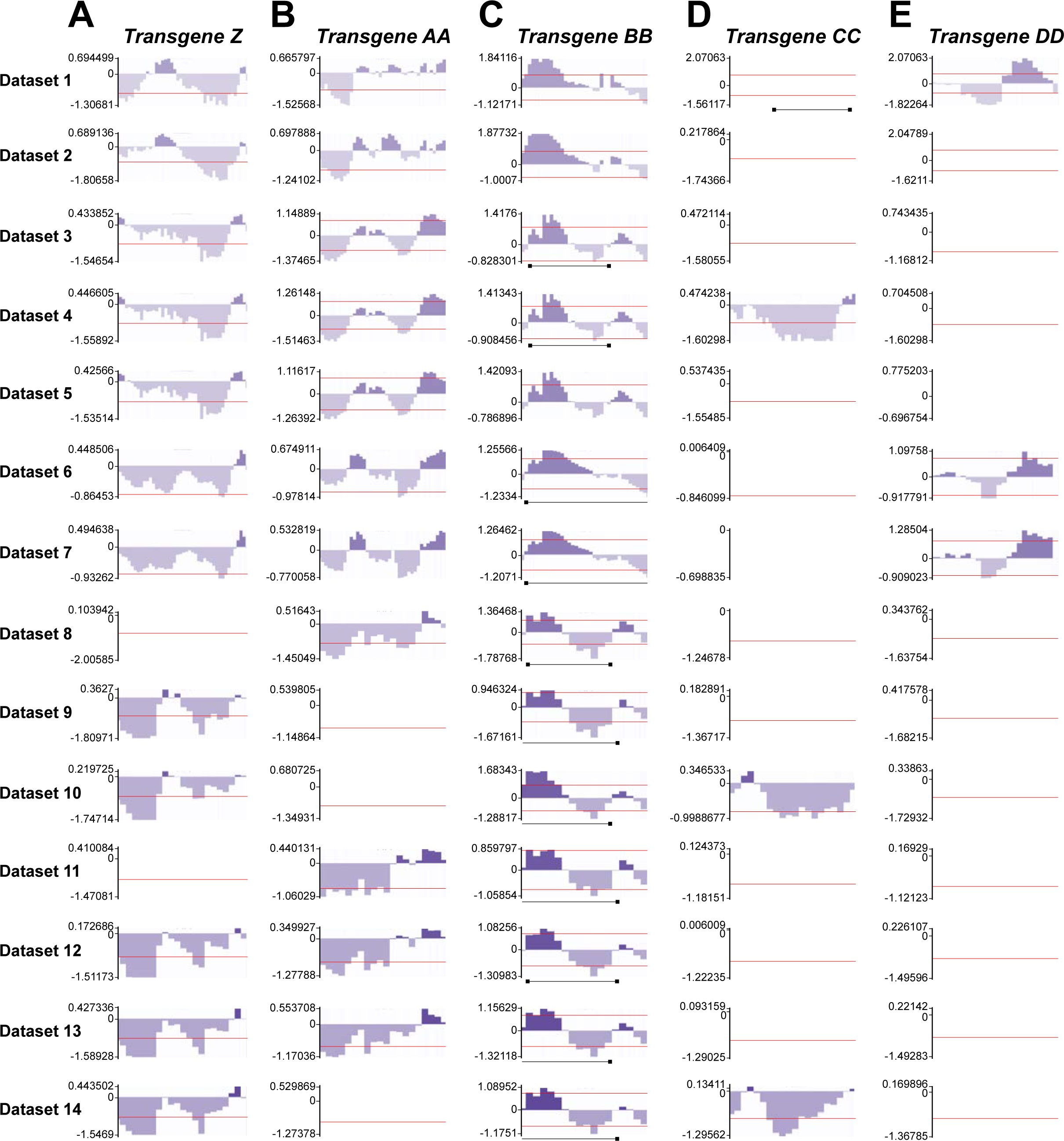
TAD calls across 14 Hi-C datasets for Transgenes Z-DD. **A-E**. Red lines indicate a directionality index signal of 0.8 or −0.8, the cutoff for a TAD. Black bars indicate TAD calls. Directionality indices are shown for the entire region spanned by each transgene.

**Supplemental Figure 8:**
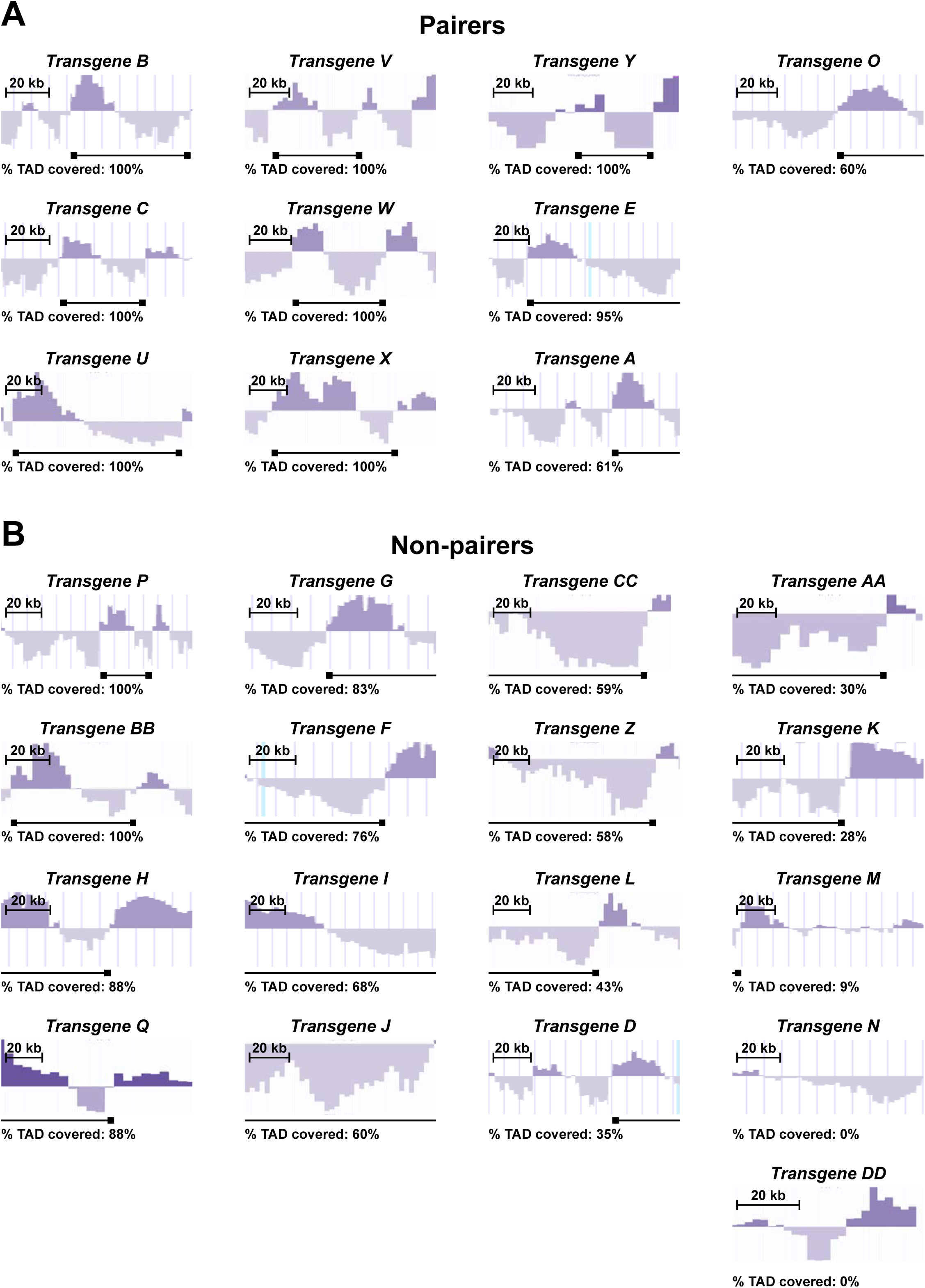
A higher percentage of pairers encompass entire TADs than non-pairers. **A**. Representative directionality indices showing the percentage of a TAD covered by each pairing transgene. Black bars indicate consensus TAD calls generated from analysis of 14 Hi-C datasets(Hou et al., 2012; Li et al., 2015; Schuettengruber et al., 2014; Stadler et al., 2017). Representative directionality indices are from NCBI accession numbers GSE38468 (*Transgenes C, U)*, GSE61471(*Transgenes A, B, E, O, W, X*), GSM2679637 (*Transgene Y*), and GSE63515 (*Transgene V*).Directionality indices are shown for the entire region spanned by each transgene. **B**. Representative directionality indices showing the percentage of a TAD covered by each non-pairing transgene. Black bars indicate consensus TAD calls generated from analysis of 14 Hi-Cdatasets(Hou et al., 2012; Li et al., 2015; Schuettengruber et al., 2014; Stadler et al., 2017).Representative directionality indices are from NCBI accession numbers GSE38468 (*Transgenes G,H, N, p*), GSE61471 (*Transgenes D, F, I, J, K, L, M, Z, BB, CC*), GSE63515 (*Transgene DD*),GSM2679644 (*Transgene Q*), and GSM2679637 (*Transgene AA*). Directionality indices are shown forthe entire region spanned by each transgene.

**Supplemental Figure 9:**
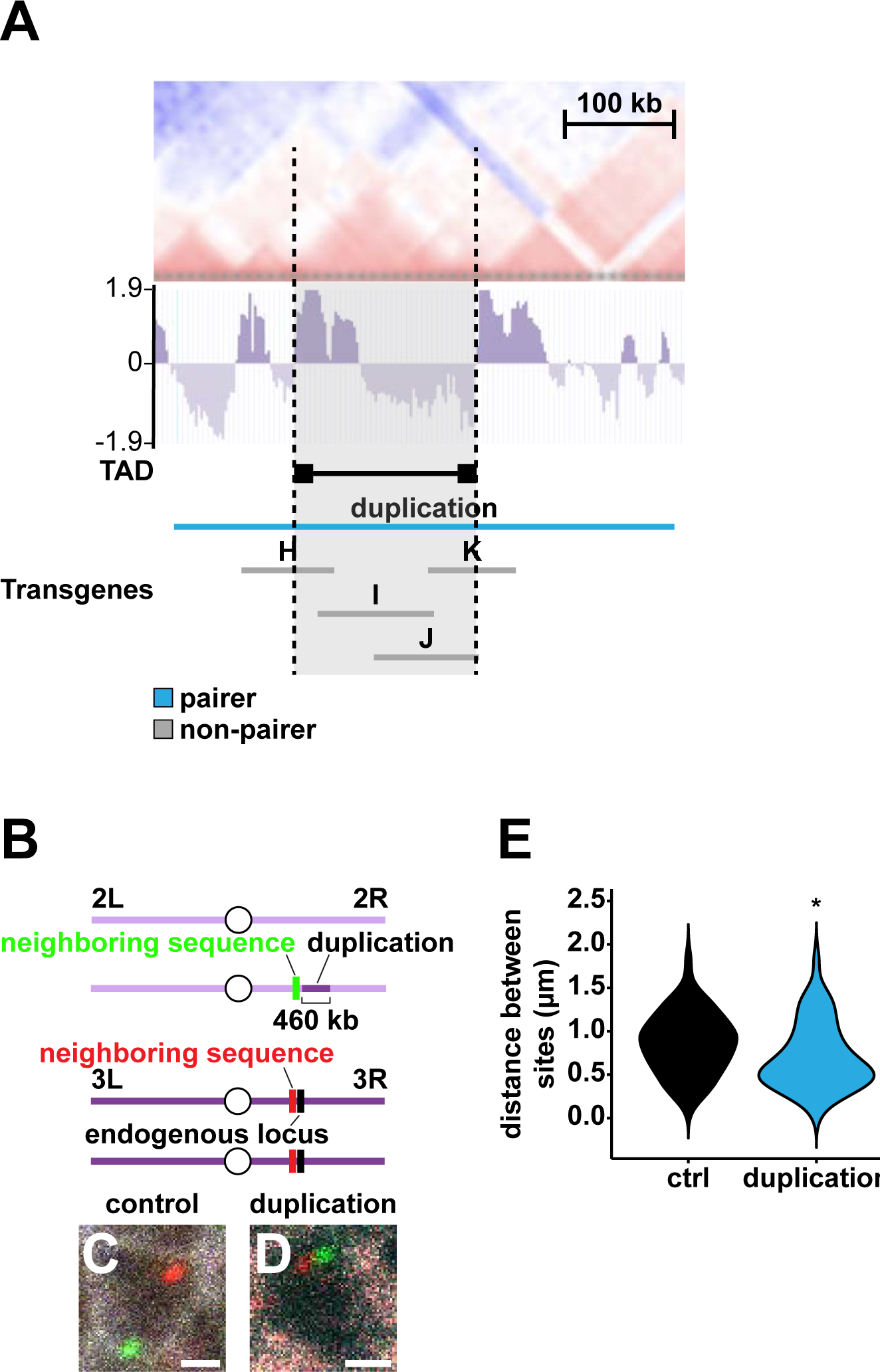
A 460-kb duplication encompassing a TAD drives pairing. **A**. Representative HiC heat map and directionality index (NCBI GSE38468) showing large TAD covered by the duplication. Dotted lines: TAD boundaries. Black bar: TAD. See **Fig. S5A** for TADassessment. **B**. DNA FISH strategy used to assess pairing of the duplication with its endogenous locus. **C-D**. Control and duplication. Scale bars=1 μm. White: Lamin B, red: probes neighboring endogenous locus, green: probes neighboring duplication breakpoint. **E**. Quantifications for **Fig. S9C-D**. Black: control, blue: pairer. *=p<0.05, Wilcoxon rank-sum test.

**Supplemental Figure 10:**
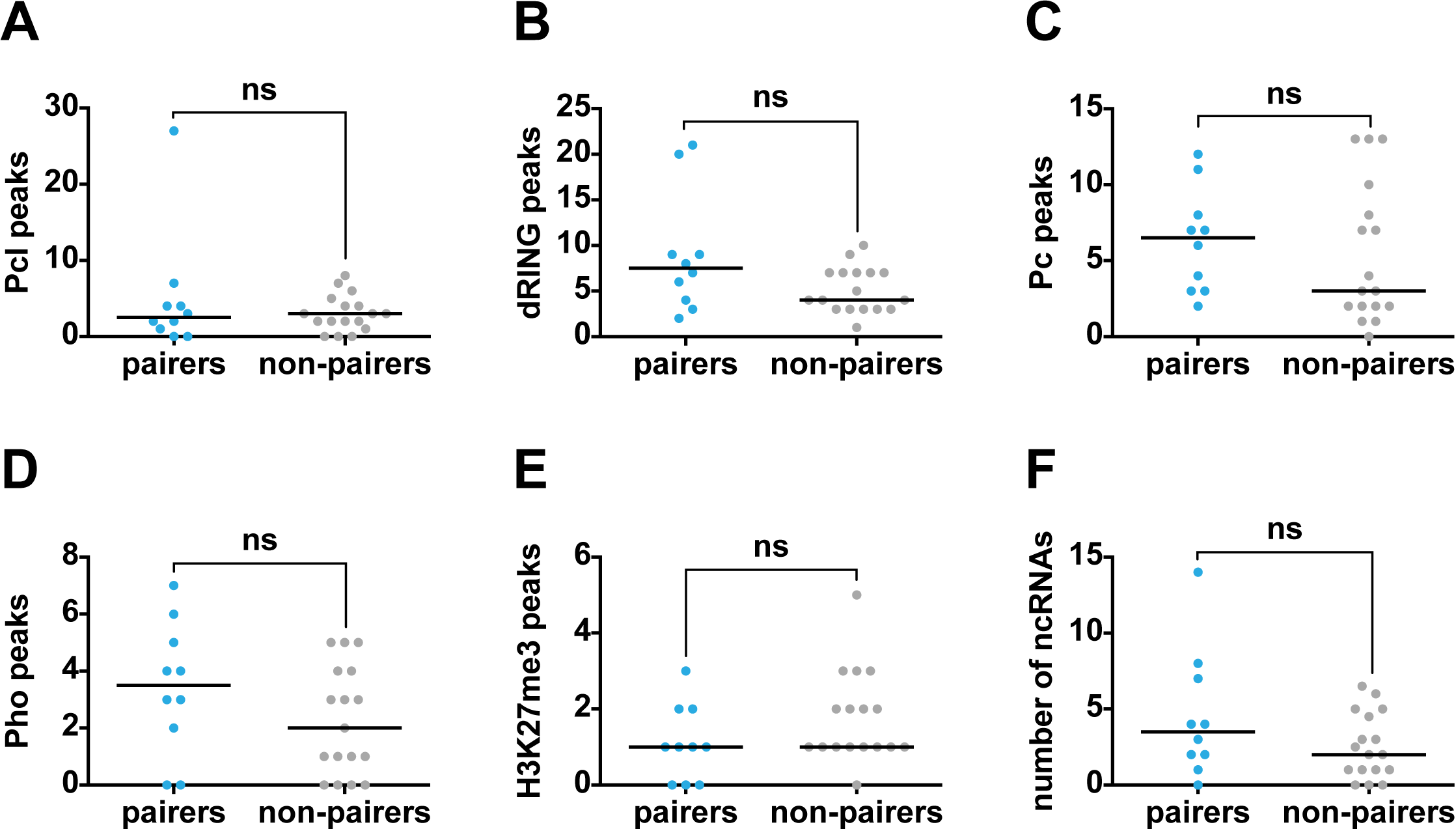
Polycomb Group Complex binding sites, repressive chromatin marks, and ncRNAs do not account for pairing. **A-F**. Graphs showing the number of Polycomb Group Complex or H3K27me3 ChIP peaks or the number of ncRNAs per transgene for all pairers and non-pairers tested in **Fig. 2F**, **3E**, and **S2E**. Blue: pairers, gray: non-pairers. ns=p>0.05, Wilcoxon rank-sum test. Black lines indicate medians.

**Supplemental Figure 11:**
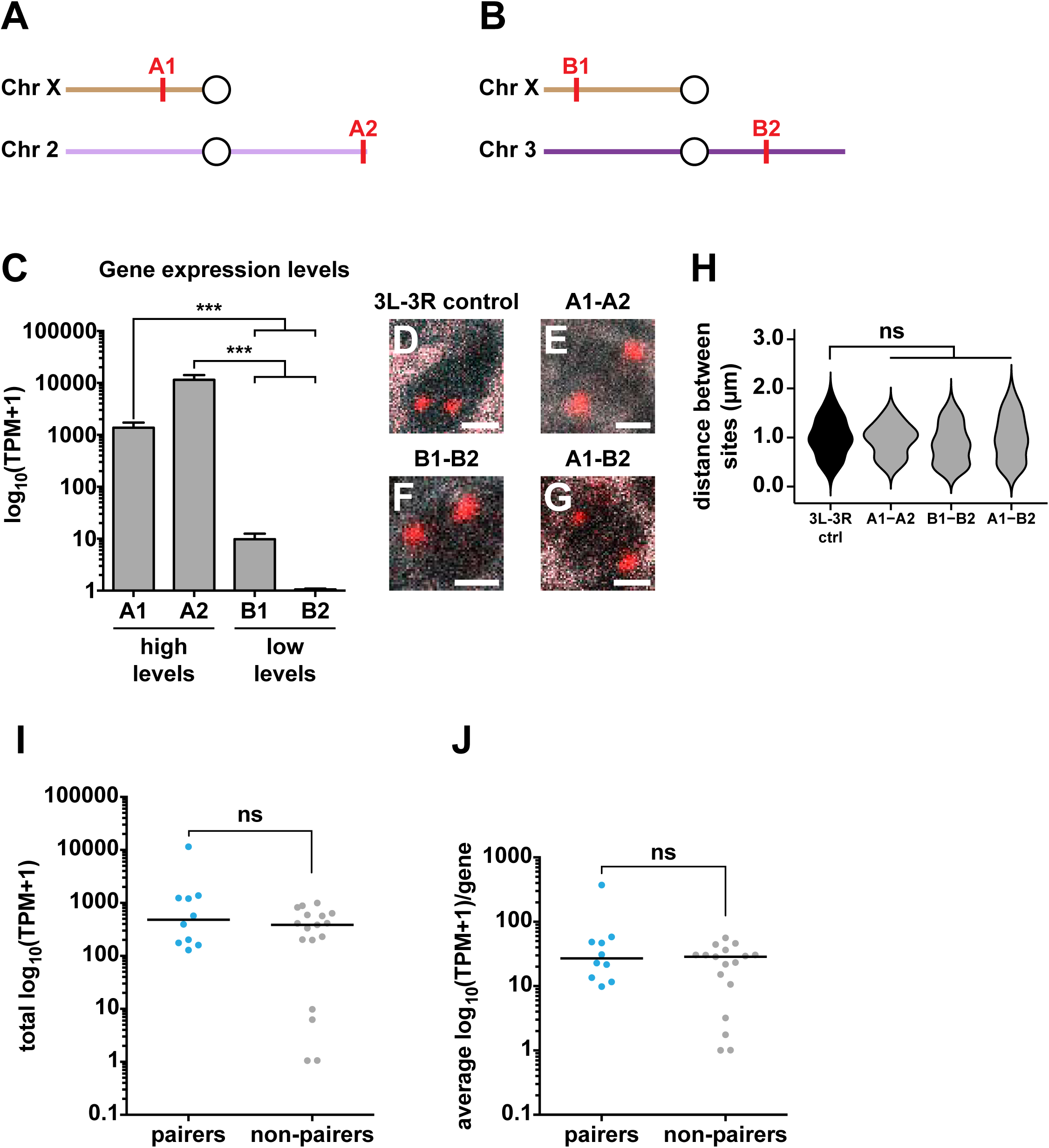
Compartmentalization and gene expression state do not account for pairing. **A-B**. Schematics indicating genomic locations of high-expressing loci A1 and A2 (**A**) and low-expressing loci B1 and B2 (**B**). **C**. TPM+1 values for A and B compartment regions tested for interactions. **D-G**. Example images for 3L-3R control, A1-A2, B1-B2, and A1-B2 compartment experiments. Scale bars=1 μm. White: Lamin B, red: probes against transgene insertion site and endogenous site (**D**) or two compartment sites (**E-G**). 3L-3R control image is from the same experiment as in **Fig. S4B** and **S14D**. **H**. Quantifications for **S11D-G**. Black: control, gray: non-pairers. ns=p>0.05, one way ANOVA on ranks with Dunn’s multiple comparisons test. 3L-3R control data are the same as in **Fig. S1A** (3L-3R control), **S4D**, and **S14Q** (3L-3R control). **I-J**. ns=p>0.05, Wilcoxon rank-sum test. Black lines indicate medians. **I**. Total TPM+1 per transgene. **J**. Average TPM+1/gene for each transgene.

**Supplemental Figure 12:**
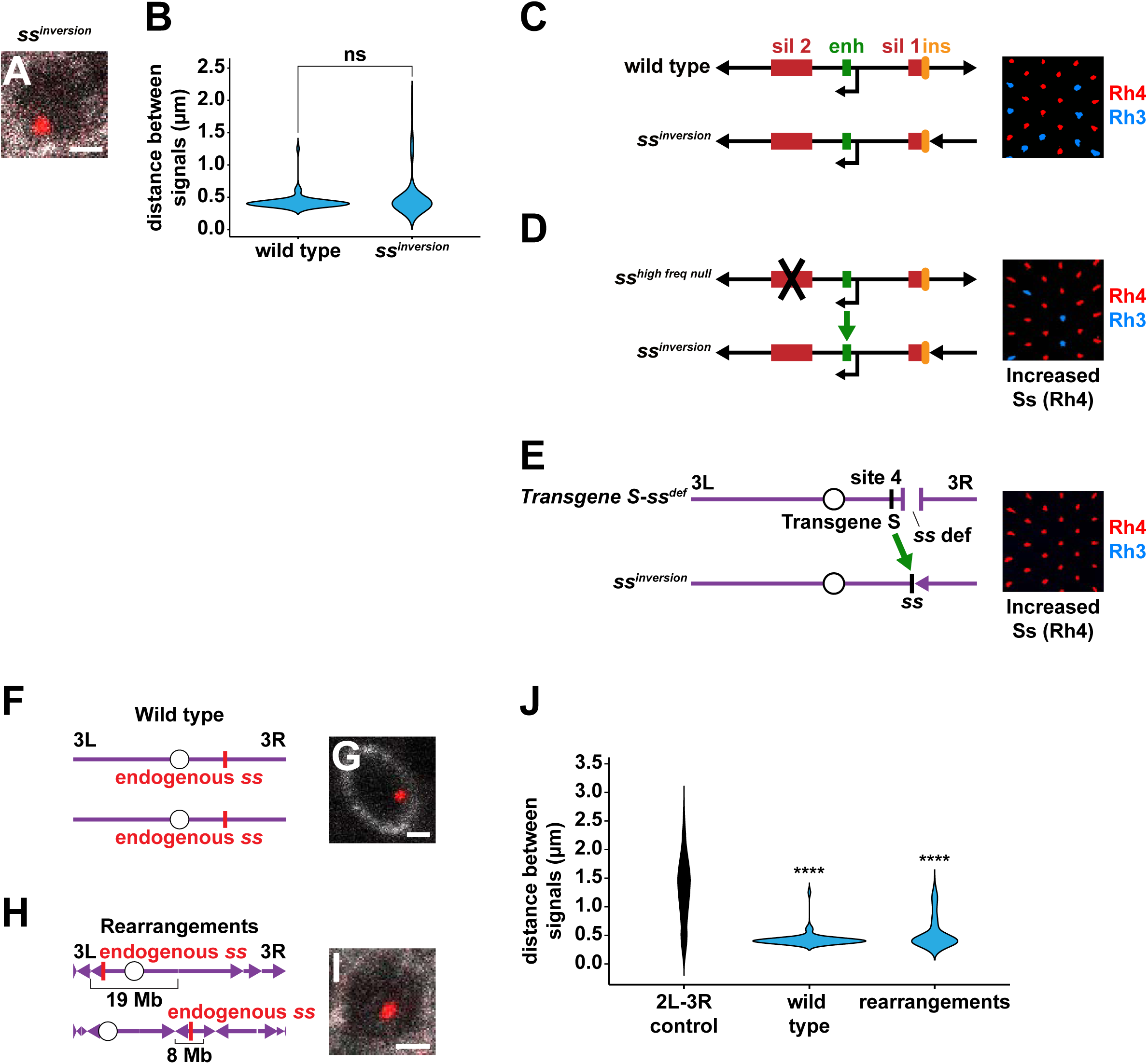
The *ss* button drives pairing and transvection despite chromosome rearrangements. **A**. *ss*^*inversion*^ / + example. Scale bar=1 μm. White: Lamin B, red: probes against endogenous *ss*. **B**. Quantification of **Fig. S12A**. ns=p>0.05, Wilcoxon rank-sum test. Data for wild type are the sameas in **Fig. S12J**. Blue: pairers. **C**. Schematic and representative image of *ss*^*inversion*^ / + adult R7s. Ss(Rh4)=63%. *ss*^*inversion*^ / + had noeffect on the normal Rh3:Rh4 ratio. ins: insulator, sil 1: silencer 1, enh: enhancer, sil 2: silencer 2.Smaller black arrows: transcription start sites. Red: Rh4, blue: Rh3. **D**. Schematic and representative image of *ss*^*inversion*^ / *ss*^*high freq null*^ adult R7s. Ss(Rh4)=80%. *ss*^*high freq null*^produces no functional Ss protein, but it performs transvection to increase the expression frequency of *ss* on other chromosomes (Johnston and Desplan, 2014). *ss*^*high freq null*^ upregulated expressionfrequency from *ss*^*inversion*^, indicating that *ss*^*inversion*^ performed transvection. Black X indicates that there is a mutation in the second silencer of *ss* that disrupts the protein coding sequence of *ss*^*high freq null*^.Smaller black arrows: transcription start sites. Green arrow indicates upregulation of *ss* between chromosomes. Red: Rh4, blue: Rh3. **E**. Schematic and representative image of *Transgene S-ss*^*def*^ / *ss*^*inversion*^ adult R7s. Ss(Rh4)=99%.*Transgene S* was recombined onto a chromosome with a *ss* deficiency to examine *Transgene S*transvection with mutant *ss* alleles. *Transgene S* performs transvection to upregulate expression of wild type, endogenous *ss* (Johnston and Desplan, 2014). *Transgene S* upregulated expression of *ss*on the *ss*^*inversion*^ allele, indicating that *ss*^*inversion*^ performed transvection. Red: Rh4, blue: Rh3. Green arrow indicates upregulation of *ss* between chromosomes. **F&H**. DNA FISH strategies used to assess endogenous *ss* pairing in wild type and chromosome rearrangement backgrounds. **G&I**. Wild type and chromosome rearrangement examples. Scale bars=1 μm. White: pm181 (R7 marker)>GFP in **G**, Lamin B in **I**. Red: probes against endogenous *ss.* **J**. Quantifications for **Fig. S12G, I**. ****=p<0.0001, one-way ANOVA on ranks with Dunn’s multiple comparisons test. 2L-3R control data are the same as in **Fig. 2F, 3E** (2L-3R control), **3I, 6C** (2L-3R eye control), **S1A-B, S14Q** (2L-3R control). Data for wild type are the same as in **Fig. S11B**. Black: control, blue: pairers.

**Supplemental Figure 13:**
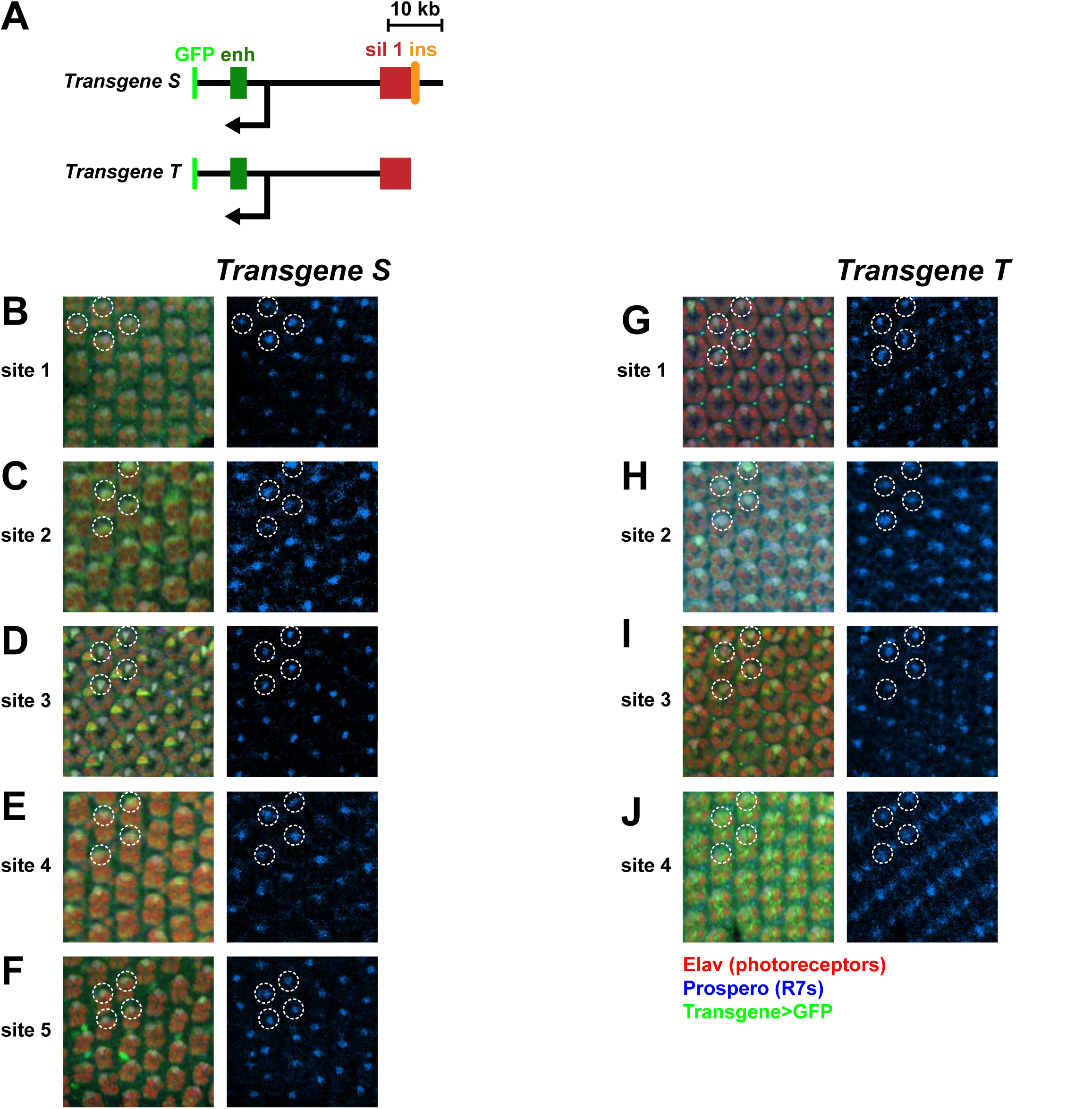
*Transgenes S* and *T* are expressed in 100% of R7 photoreceptors at all insertion sites. **A**. Schematic of *Transgenes S* and *T* with GFP tags. ins: insulator, sil1: silencer, enh: enhancer. Black arrows: transcription start sites. **B-F**. Representative images of *Transgene S*>GFP expression in mid-pupal R7 photoreceptors for all insertion sites. Red: Elav (photoreceptors), blue: Prospero (R7s), green: *Transgene S*>GFP. White circles indicate representative R7s. **G-J**. Representative images of *Transgene T*>GFP expression in mid-pupal R7 photoreceptors for all insertion sites. Red: Elav (photoreceptors), blue: Prospero (R7s), green: *Transgene T*>GFP. White circles indicate representative R7s.

**Supplemental Figure 14:**
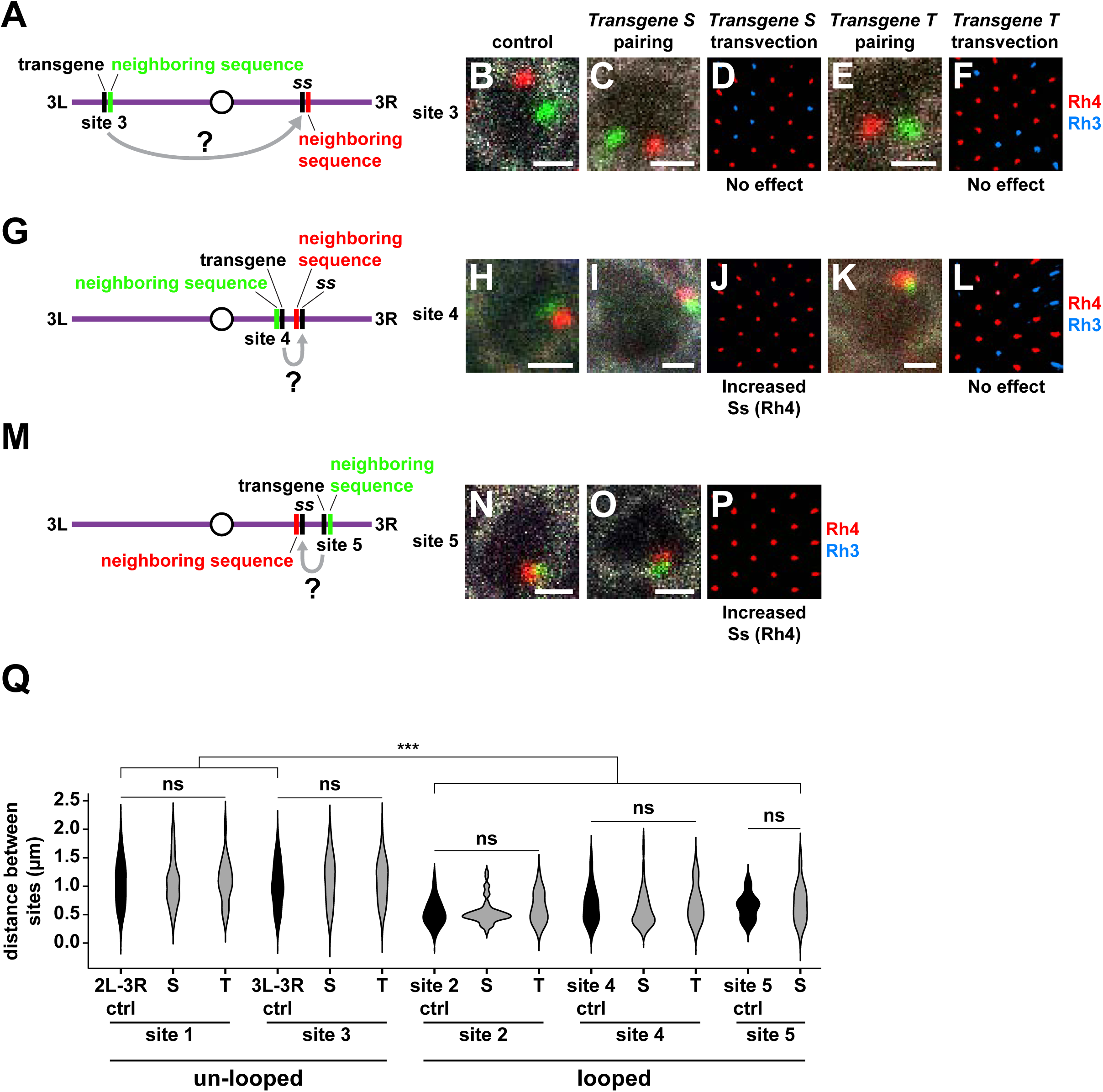
Pairing is necessary but not sufficient for *ss* transvection. **A, G, M**. DNA FISH strategies used to test pairing and transvection of *Transgenes S* and *T* at each insertion site. Gray arrow with “?” indicates that *Transgenes S* and *T* were tested for transvection. **B-C, E, H-I, K, N-O**. DNA FISH examples for control, *Transgene S*, and *Transgene T* at each insertion site. Scale bars=1 μm. White: Lamin B, red: probes neighboring endogenous sequence, green: probes neighboring transgene insertion site. Image for site 3 control is from the same experiment as in **S4B** and **S11D**. **D, F, J, L, P**. Representative images of adult eyes for *Transgene S* and *Transgene T* at each insertion site. Red: Rh4, blue: Rh3. **D**. Ss(Rh4)=71% **F**. Ss(Rh4)=76% **J**. Ss(Rh4)=98% **L**. Ss(Rh4)=74% **P**. Ss(Rh4)=98% **Q**. Quantifications for controls, *Transgene S*, and *Transgene T* at each insertion site. Black: control, gray: non-pairer. ***=p<0.0005, ns=p>0.05, ordinary one-way ANOVA with Dunnett’s multiple comparisons test (*Transgenes S* and *T site 1* vs. *2L-3R ctrl*, *Transgenes S* and *T site 3* vs. *3L-3R ctrl*), one-way ANOVA on ranks with Dunn’s multiple comparisons test (*Transgenes S* and *T site 2* vs. *site 2 control*; *Transgenes S* and *T site 4* vs. *site 4 control*; *site 2 control, Transgenes S* and *T site 2* vs. *2L-3R control* and vs. *3L-3R control*; *site 4 control, Transgene S* and *T site 4* vs. *2L-3R control* and *3L-3R control*; *site 5 control* and *Transgene S site 5* vs. *2L-3R control* and *3L-3R control*), or unpaired t-test with Welch’s correction (*Transgene S site 5*). Data for 2L-3R control are the same as in **Fig. 2F, 3E** (2L-3R control), **3I, 6C** (2L-3R eye control), **S1A-B**, and **S12J**. Data for site 3 control are the same as in **Fig. S4D** and **S11H**. Data for *Transgene S site 1* are the same as in **Fig. 3I**.

**Supplemental Figure 15:**
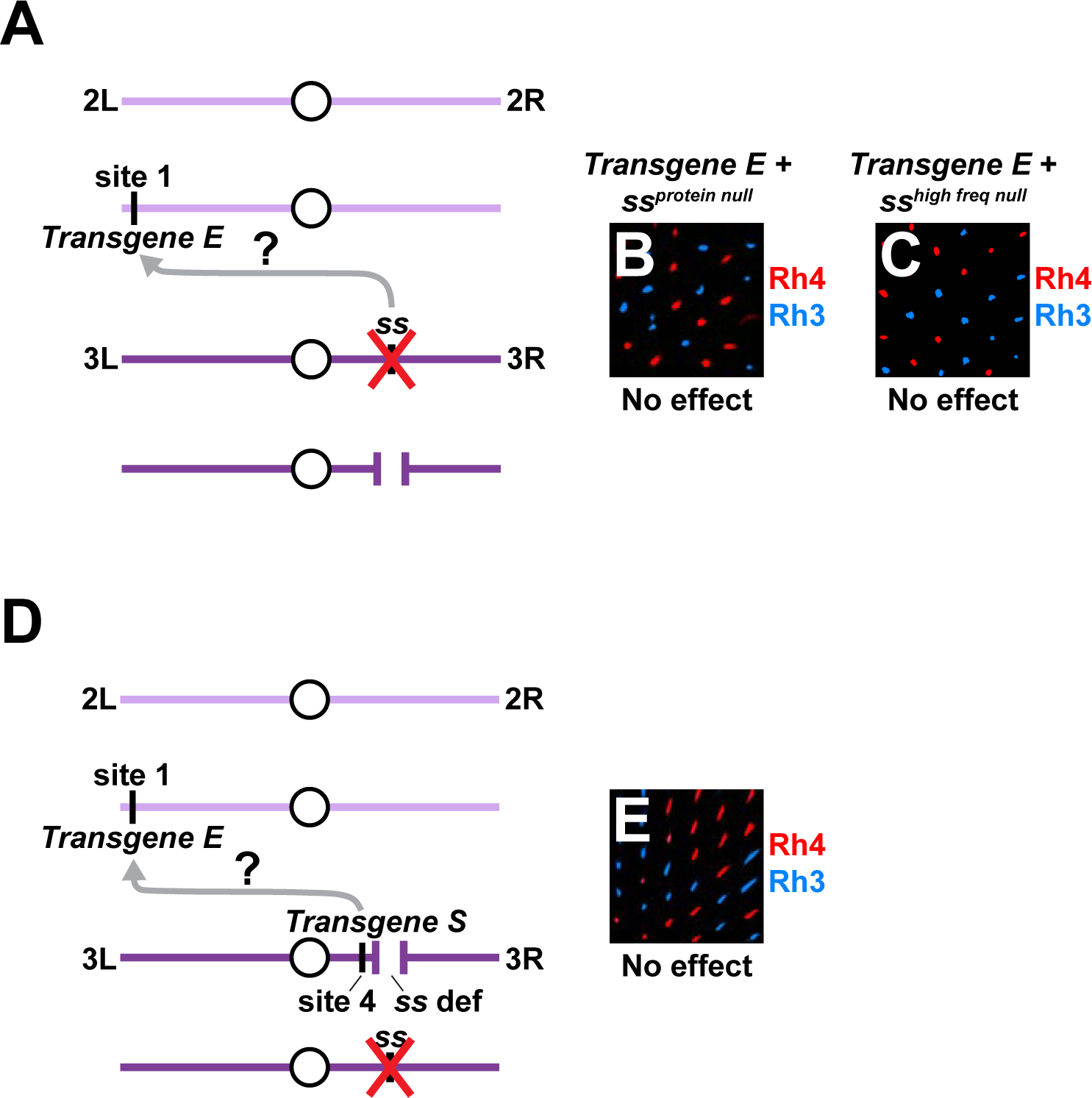
*Transgene E* does not perform transvection. **A**. Transvection assay to test whether mutant *ss* alleles alter expression from *Transgene E.* Red X indicates *ss* mutant allele, which is *ss*^*protein null*^ from **Fig. S15B** or *ss*^*high freq null*^ from **Fig. S15C**. Gray arrow with “?” indicates that *Transgene E* was tested for transvection. **B-C**. Rh3 and Rh4 expression in *Transgene E + ss*^*protein null*^ / *ss*^*def*^ (Ss(Rh4)=52%) and *Transgene E + ss*^*high freq null*^ / *ss*^*def*^ (Ss(Rh4)=51%). **D**. Schematic of *Transgene E + Transgene S-ss*^*def*^ / *ss*^*protein null*^ genotype. *Transgene S* was recombined onto a chromosome with a *ss* deficiency to examine *Transgene S* transvection with mutant *ss* alleles. *Transgene S* performed transvection to upregulate expression of wild type,endogenous *ss* (Johnston and Desplan, 2014). In the *Transgene E + Transgene S-ss*^*def*^ / *ss*^*protein null*^genotype, the endogenous *ss* locus was hemizygous for a protein coding null allele of *ss*, which produced no functional Ss protein. Therefore, any functional Ss protein in this genotype was produced by *Transgene E*, and an increase in Ss (Rh4) expression frequency indicated that*Transgene S* was performing transvection to upregulate Ss expression from *Transgene E*. Red Xover the *ss* locus indicates that the *ss* allele is a protein coding null. Gray arrow with “?” indicates that*Transgene E* was tested for transvection. **E**. Adult eye for the *Transgene E + Transgene S-ss*^*def*^ / *ss*^*protein null*^ genotype. *ss* expression frequency was not upregulated (Ss(Rh4)=53%), indicating that *Transgene S* did not perform transvection with*Transgene E*. Red: Rh4, blue: Rh3.

**Supplemental Figure 16:**
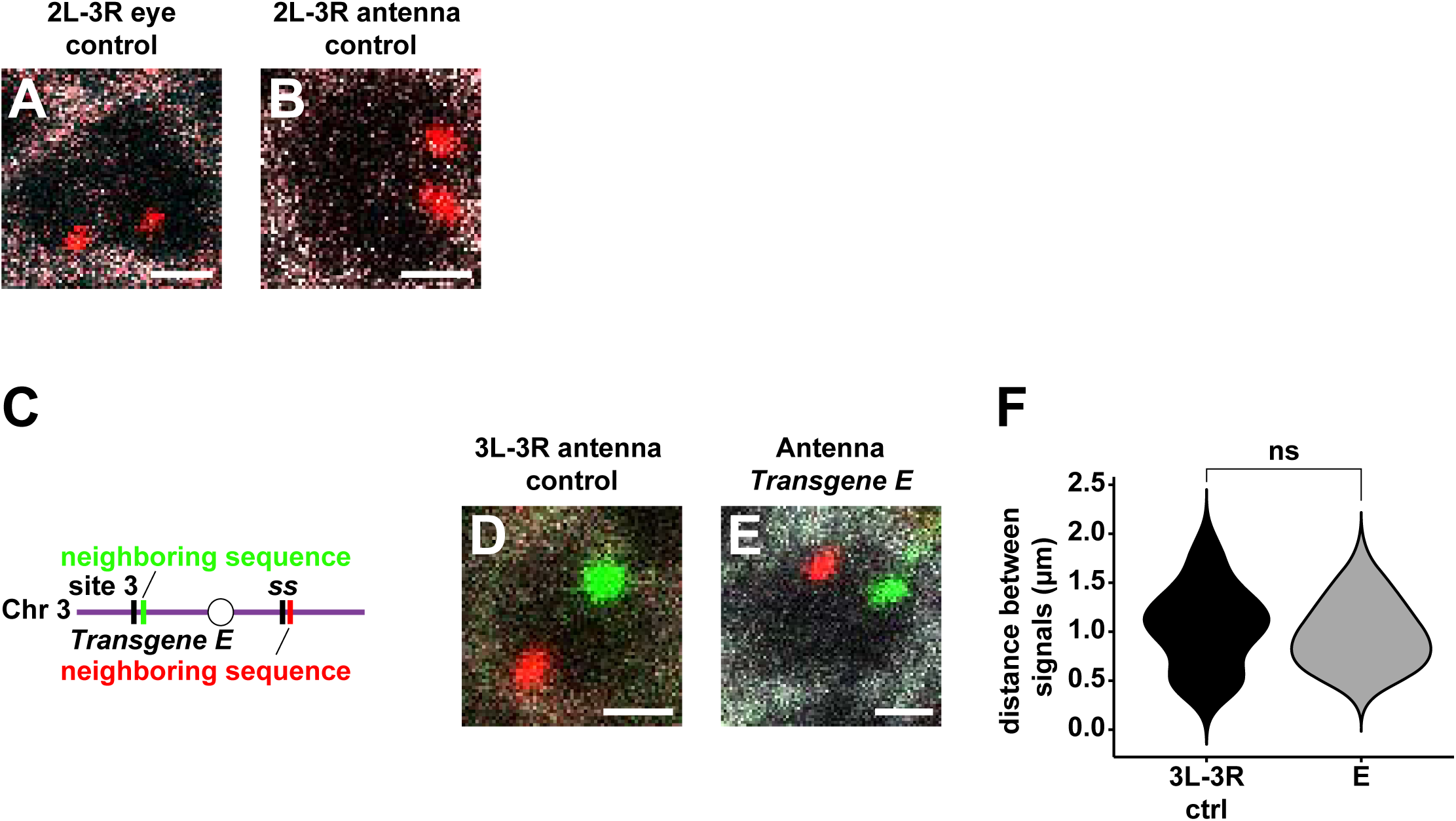
Pairing driven by the *ss* button is cell-type specific. **A-B**. *Site 1 control* images in the eye and antenna. Scale bars=1 μm. White: Lamin B, red: probes against endogenous *ss* and site 1 insertion site. Site 1 probes were pseudocolored red to allow direct comparison with single-color FISH experiments. **C**. DNA FISH strategy used to assess pairing of *Transgene E* at site 3 with its endogenous locus in the larval antenna. **D-E**. *Site 3 control* and *Transgene E site 3* examples in the larval antenna. Scale bar=1 μm. White: Lamin B, red: probes neighboring endogenous sequence, green: probes neighboring transgene insertion site. **F**. Quantifications for **Fig. S16B-C**. ns=p>0.05, unpaired t-test with Welch’s correction.

**Supplemental Figure 17:**
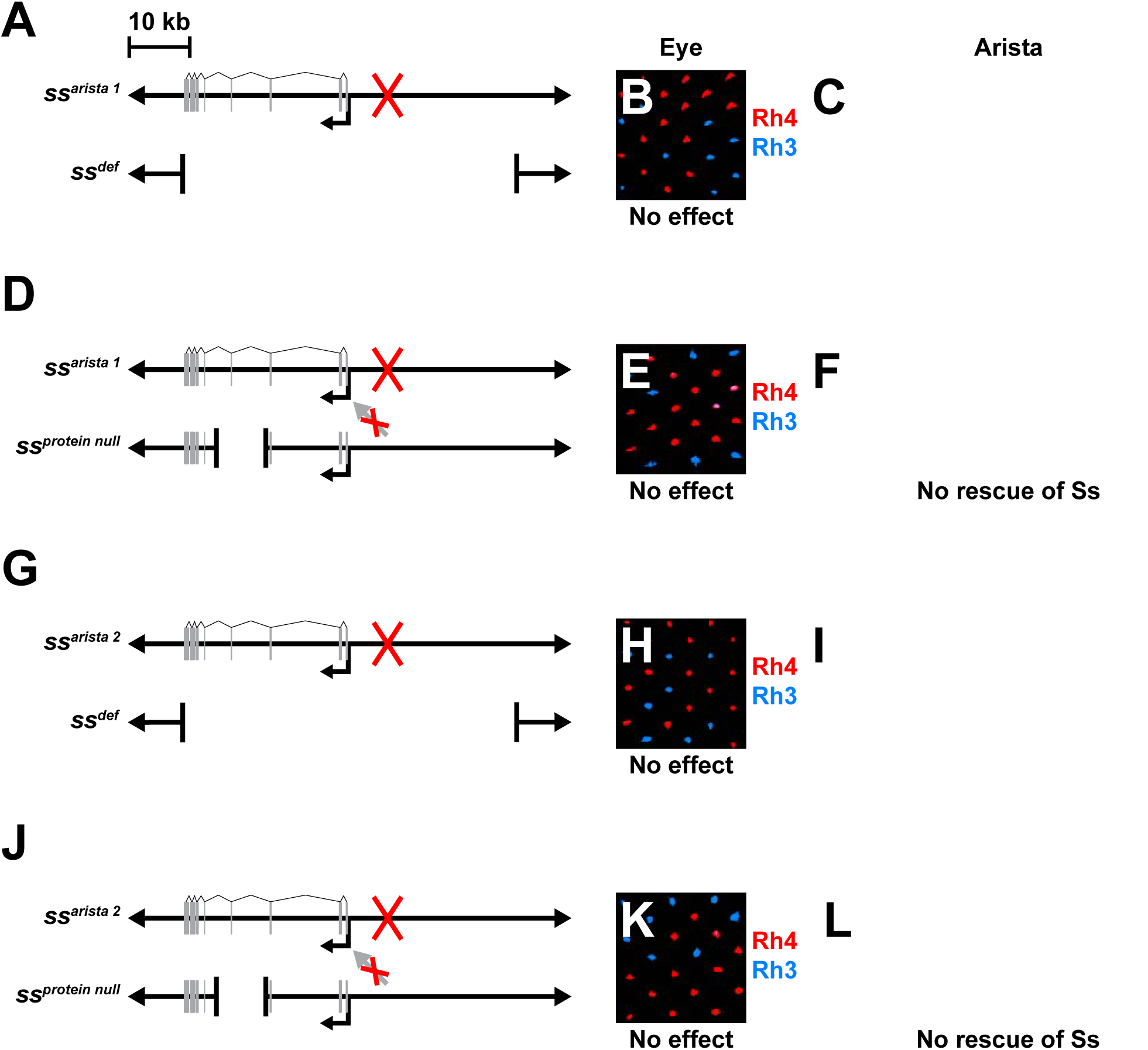
*ss* mutant alleles with arista-specific phenotypes do not perform transvection in the arista. **A, D, G, J**. Genotypes tested for transvection. Gray rectangles: exons. Smaller black arrows:transcription start sites. Red X indicates an uncharacterized mutation in the *ss*^*arista 1*^ or *ss*^*arista 2*^sequence. Red X over gray arrow indicates an absence of transvection between alleles in the arista. **B**. *ss*^*arista 1*^ / *ss*^*def*^ adult eye. *ss*^*arista 1*^ / *ss*^*def*^ had no effect on eye development. Red: Rh4, blue: Rh3.Ss(Rh4)=53%. **C**. *ss*^*arista 1*^ / *ss*^*def*^ arista. *ss*^*arista 1*^ / *ss*^*def*^ caused aristapedia. Scale bar= 50 μm. White arrow indicates arista. Image is from the same experiment as in **Fig. 6I**. **E**. *ss*^*arista 1*^ / *ss*^*protein null*^ adult eye. *ss*^*arista 1*^ / *ss*^*protein null*^ had no effect on eye development. Red: Rh4,blue: Rh3. Ss(Rh4)=53%. **F**. *ss*^*arista 1*^ / *ss*^*protein null*^ arista. *ss*^*arista 1*^ / *ss*^*protein null*^ had aristapedia, indicating that transvection did not occur to rescue the mutant *ss* phenotype. Scale bar= 50 μm. White arrow indicates arista. **H**. *ss*^*arista 2*^ / *ss*^*def*^ adult eye. *ss*^*arista 2*^ / *ss*^*def*^ had no effect on eye development. Red: Rh4, blue: Rh3. Ss(Rh4)=60%. **I**. *ss*^*arista 2*^ / *ss*^*def*^ arista. *ss*^*arista 2*^ / *ss*^*def*^ caused aristapedia. Scale bar= 50 μm. White arrow indicates arista. **K**. *ss*^*arista 2*^ / *ss*^*protein null*^ adult eye. *ss*^*arista 2*^ / *ss*^*protein null*^ had no effect on eye development. Red: Rh4, blue: Rh3. Ss(Rh4)=62%. **L**. *ss*^*arista 2*^ / *ss*^*protein null*^ arista. *ss*^*arista 2*^ / *ss*^*protein null*^ had aristapedia, indicating that transvection did not occur to rescue the mutant *ss* phenotype. Scale bar= 50 μm. White arrow indicates arista.

**Supplemental Figure 18:**
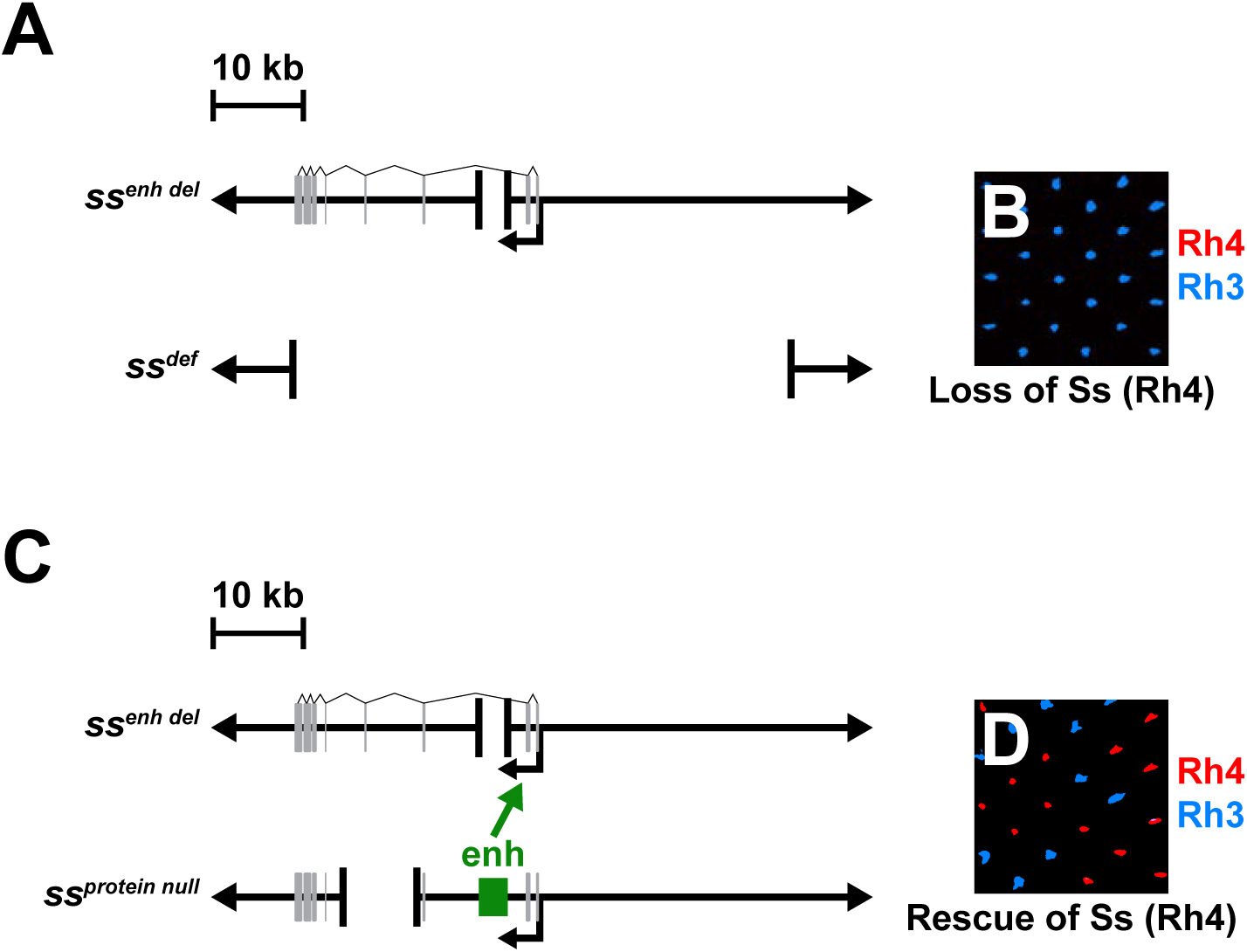
*ss*^*protein null*^ performs transvection in the eye. **A**. Schematic of *ss* enhancer deletion (*ss*^*enh del*^) allele over a *ss* deficiency (*ss*^*def*^). Gray rectangles: exons. Smaller black arrow: transcription start site. **B**. *ss*^*enh del*^ / *ss*^*def*^ adult eye. *ss*^*enh del*^ / *ss*^*def*^ caused a near complete loss of Ss/Rh4 expression. Red:Rh4, blue: Rh3. Ss(Rh4)=0.1%. **C**. Schematic of *ss*^*enh del*^ over *ss*^*protein null*^. Through transvection, the functional enhancer of *ss*^*protein null*^acted on the functional protein coding region of *ss*^*enh del*^ to rescue Ss expression (green arrow). Enh:enhancer, gray rectangles: exons. Smaller black arrows: transcription start sites. **D**. *ss*^*enh del*^ / *ss*^*protein null*^ adult eye. *ss*^*protein null*^ rescued Ss expression from the *ss*^*enh del*^ allele. Red: Rh4,blue: Rh3. Ss(Rh4)=59%.

**Supplemental Figure 19:**
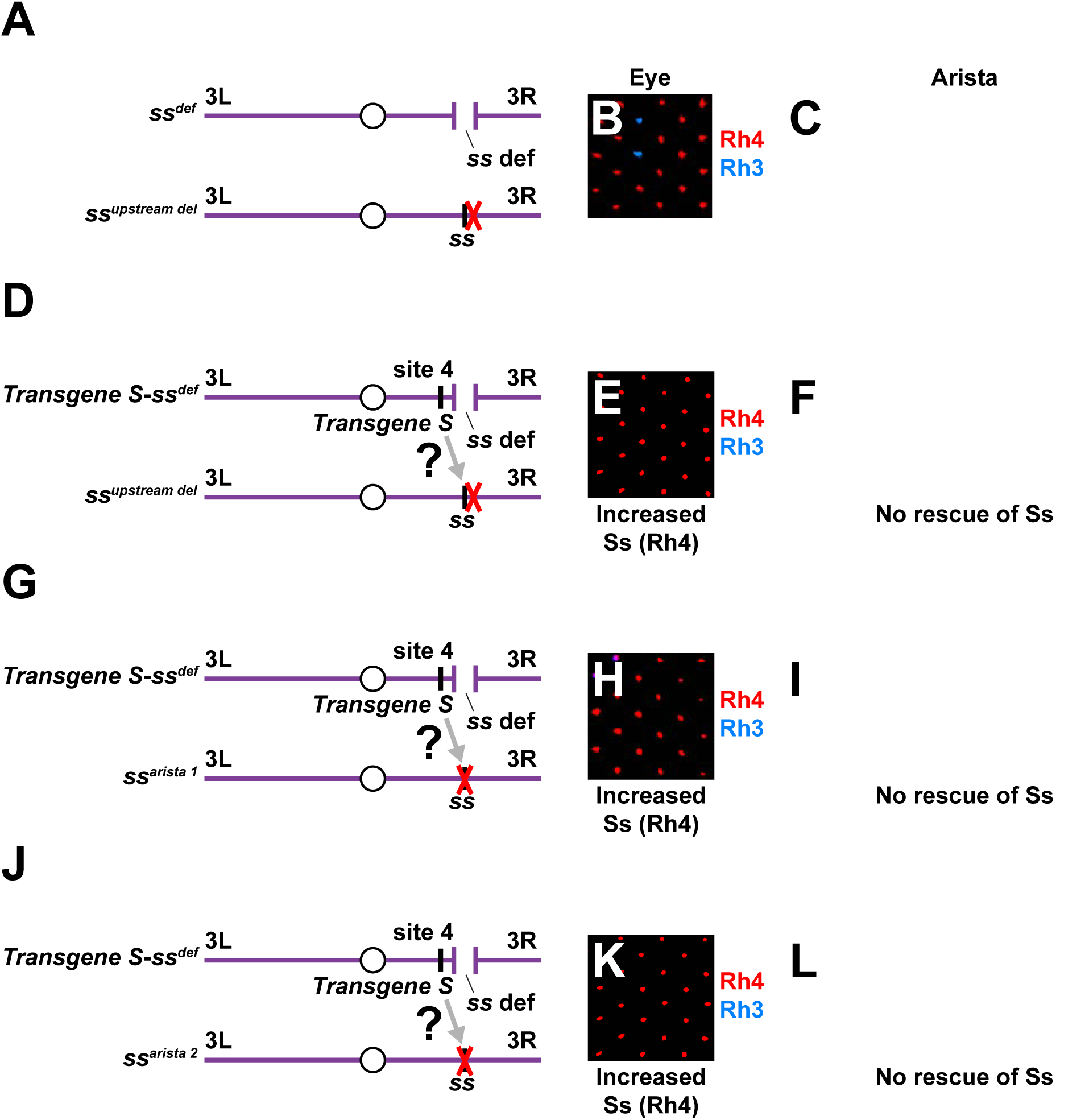
Transgene S performs transvection in the eye but not in the arista. **A**. Schematic of the *ss*^*upstream del*^ allele over the *ss*^*def*^ allele. *ss*^*upstream del*^ is a CRISPR allele in which12.7 kb of the upstream regulatory regions of *ss* are deleted. Red X indicates the deletion of regulatory regions directly upstream of the *ss* locus. **B**. *ss*^*upstream del*^ / *ss*^*def*^ adult eye. *ss*^*upstream del*^ displayed Ss(Rh4) expression in 85% of R7s. Red: Rh4,blue: Rh3. **C**. *ss*^*upstream del*^ / *ss*^*def*^ arista. *ss*^*upstream del*^ caused aristapedia. Scale bar= 50 μm. White arrow indicates arista. **D**. Schematic of the *Transgene S-ss*^*def*^ allele over the *ss*^*upstream del*^ allele. *Transgene S* was recombined onto a chromosome with a *ss* deficiency to examine *Transgene S* transvection with mutant *ss* alleles.Red X indicates the deletion of regulatory regions directly upstream of the *ss* locus. Gray arrow with a“?” indicates that *Transgene S* was tested for transvection with the *ss*^*upstream del*^ allele. **E**. *Transgene S-ss*^*def*^ / *ss*^*upstream del*^ adult eye. *Transgene S-ss*^*def*^ upregulated Ss(Rh4) expression fromthe *ss*^*upstream del*^ allele into 100% of R7s, indicating that *Transgene S* performed transvection with *ss*^*upstream del*^ in the eye. Red: Rh4, blue: Rh3. **F**. *Transgene S-ss*^*def*^ / *ss*^*upstream del*^ arista. *Transgene S-ss*^*def*^ / *ss*^*upstream del*^ had aristapedia, indicating that *Transgene S* did not perform transvection to rescue *ss*^*upstream del*^ expression in the arista. Scalebar= 50 μm. White arrow indicates arista. **G**. Schematic of the *Transgene S-ss*^*def*^ allele over the *ss*^*arista 1*^ allele. Red X indicates an uncharacterized mutation in the *ss*^*arista 1*^ allele. Gray arrow with “?” indicates that *Transgene S* was tested for transvection with the *ss*^*arista 1*^ allele. **H**. *Transgene S-ss*^*def*^ / *ss*^*arista 1*^ adult eye. *Transgene S-ss*^*def*^ upregulated Ss(Rh4) expression from the *ss*^*arista 1*^ allele into 99% of R7s, indicating that *Transgene S* performed transvection with *ss*^*arista 1*^ in the eye. Red: Rh4, blue: Rh3. **I**. *Transgene S-ss*^*def*^ / *ss*^*arista 1*^ arista. *Transgene S-ss*^*def*^ / *ss*^*arista 1*^ had aristapedia, indicating that *Transgene S* did not perform transvection to rescue *ss*^*arista 1*^ expression in the arista. Scale bar= 50 μm. White arrow indicates arista. **J**. Schematic of the *Transgene S-ss*^*def*^ allele over the *ss*^*arista 2*^ allele. Red X indicates an uncharacterized mutation in the *ss*^*arista 2*^ allele. Gray arrow with “?” indicates that *Transgene S* was tested for transvection with the *ss*^*arista 2*^ allele. **K**. *Transgene S-ss*^*def*^ / *ss*^*arista 2*^ adult eye. *Transgene S-ss*^*def*^ upregulated Ss(Rh4) expression from the *ss*^*arista 2*^ allele into 100% of R7s, indicating that *Transgene S* performed transvection with *ss*^*arista 2*^ in the eye. Red: Rh4, blue: Rh3. **L**. *Transgene S-ss*^*def*^ / *ss*^*arista 2*^ arista. *Transgene S-ss*^*def*^ / *ss*^*arista 2*^ had aristapedia, indicating that *Transgene S* did not perform transvection to rescue *ss*^*arista 2*^ expression in the arista. Scale bar= 50 μm. White arrow indicates arista.

**Supplemental Figure 20:**
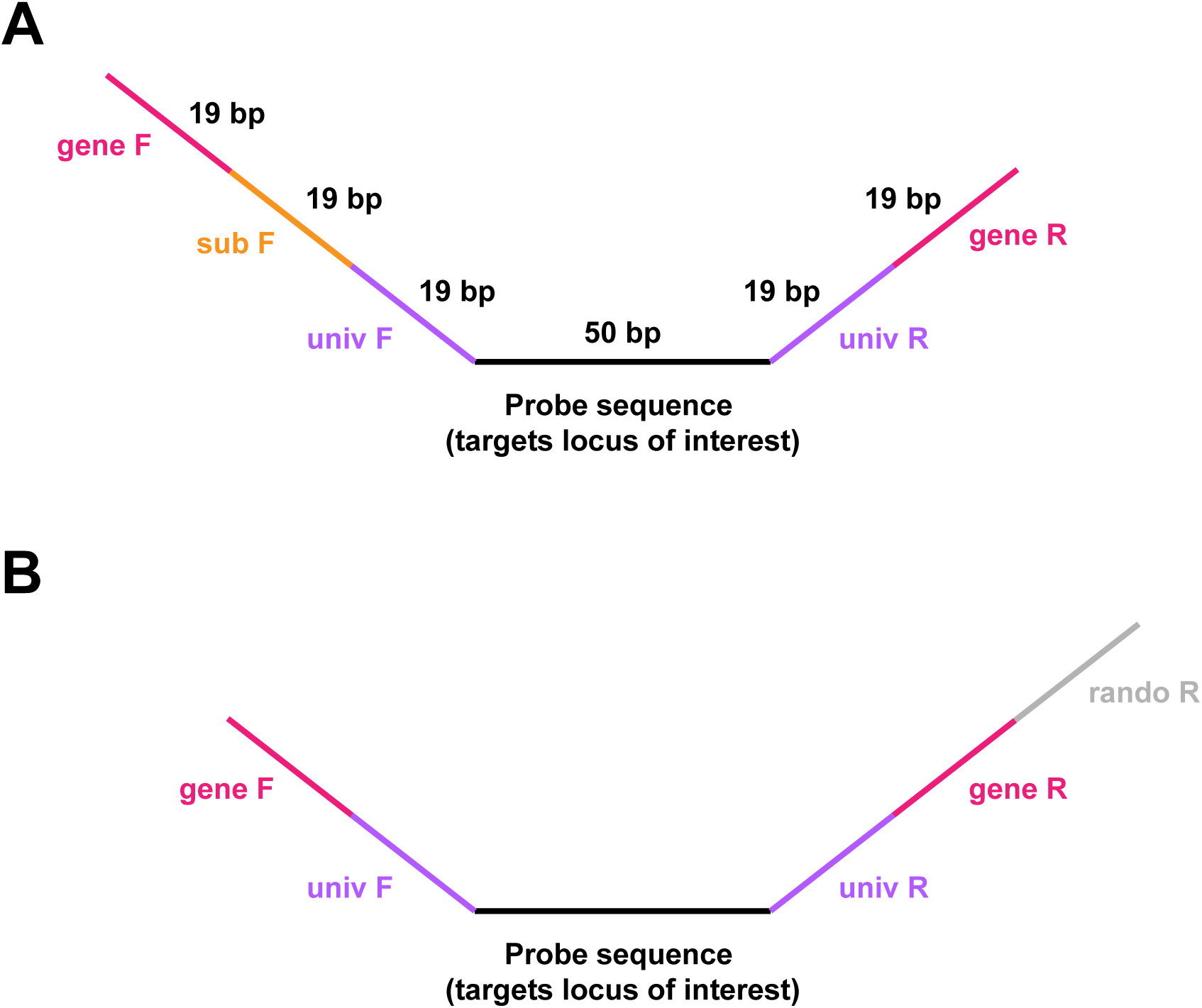
Barcoding primer scheme for DNA Oligopaints FISH probes. **A**. Schematic of barcoding primer scheme for Oligopaints probe libraries containing sublibraries. univ:universal primer, sub: sublibrary primer. **B**. Schematic of barcoding primer scheme for Oligopaints probe libraries without sublibraries. univ:universal primer, rando: random primer.

